# Disrupted priming within draining lymph nodes drives immune quiescence in gastric cancer

**DOI:** 10.1101/2025.05.05.651897

**Authors:** Sohrab Salehi, Emily E. Stroobant, Hannah Lees, Ya-Hui Lin, Shoji Shimada, Miseker Abate, Matthew J. Zatzman, Nicholas Ceglia, Samuel S. Freeman, Monika Laszkowska, Steven Maron, Andrew McPherson, Nicole Rusk, Eliyahu Havasov, Harrison Drebin, Ping Gu, Laura H. Tang, Yelena Y. Janjigian, Ruslan Soldatov, Ronan Chaligne, Sohrab P. Shah, Vivian E. Strong, Santosha A. Vardhana

## Abstract

The gastric mucosa is characterized by continuous innate immune surveillance and inflammatory signaling, yet a high proportion of gastric carcinomas (GCs) are recalcitrant to immune-directed therapies. The mechanisms by which GCs evade adaptive immune surveillance within the highly antigenic microenvironment of the gastric mucosa remains unknown. To address this, we collected patient-matched tumor tissue, distant normal tissue, metastasis, and draining lymph nodes to generate a large-scale single-cell immune profiling dataset from 64 patients (n=179 samples, >150,000 cells). From single cell analysis, we identified two distinct sources of impaired tumor surveillance within tumor draining lymph nodes. First, we observed that a significant fraction of tumor draining lymph nodes had undergone cytokine-driven reprogramming, leading to reduced dendritic cell homing and limited T cell priming. Second, T cells undergoing successful activation exhibited limited expansion and constrained differentiation, marked by expression of the quiescence-associated transcription factor Kruppel-like Factor 2 (*KLF2*). Overexpression of *KLF2* in primary T cells limited both their differentiation and cytotoxic capacity. These findings implicate both impaired T cell priming and *KLF2*-dependent T cell quiescence in limiting T cell immunity in gastric adenocarcinoma. We suggest these findings represent an emerging model for immune silencing in tumors developing from tissues with chronic inflammation.

## Introduction

Tumor growth and progression require evasion of immune surveillance; yet paradoxically, chronic inflammation is a well-established risk factor for multiple cancers. This dichotomy is evident in gastric adenocarcinoma, a malignancy with rising global incidence and poor outcomes, particularly once lymphatic or distant spread occurs^1^. The role of chronic inflammation in gastric carcinogenesis was first established with the classification of *Helicobacter pylori* as a carcinogen in 1994^2^. Subsequent associations with microbiota and autoimmune conditions further positioned chronic inflammation as a critical promotion factor of gastric carcinogenesis^3–6^. As most gastric cancers, and particularly those without evidence of microsatellite instability, are immunologically silent with sparse infiltration of cytotoxic CD8+ T lymphocytes (TILs)^7^, we sought to determine if chronic inflammation disrupts CD8+ T cell-dependent immune surveillance. We recently found that intra-tumoral T cell abundance in gastric cancer correlates with effective T cell priming in tumor-draining lymph nodes^8^. To test if chronic inflammation impairs adaptive immune surveillance by disrupting T cell priming within draining lymph nodes, we developed a surgical collection protocol enabling paired profiling of immune cells from gastric tumors and matched tumor-draining lymph nodes. We then performed an integrated analysis of single-cell RNA sequencing, T-cell receptor (TCR) sequencing and cell-surface epitope assessment (CITE-Seq)^9^ on the matched samples and validated the resultant phenotypes with functional assays. Our findings establish two complementary mechanisms by which sustained innate inflammation disrupts anti-tumor T cell immunity: by i) reducing the frequency of successful dendritic cell-T cell priming events within draining lymph nodes and ii) stabilizing transcriptional programs that reinforce T cell quiescence.

### Patient cohort, sample collection and single cell data generation

To characterize adaptive immune networks during both the priming and effector stages of the immune response, we developed a tissue collection protocol from patients with gastric adenocarcinoma, emphasizing tumor resections paired with matched draining lymph nodes (dLN) (**Fig. 1a**). 64 patients with gastric adenocarcinoma were included, broadly distributed across gastric cancer stages, subtypes and established covariates (**Extended Data Table 1, 2**). This included at least eight patients from each tumor subtype by Lauren classification^10^, and representation from all TCGA subtypes^11^ (**Fig. 1b**). Importantly, PD-L1 expression by combined positive score (CPS)^12^ spanned the full distribution (0-100), indicating representation across a broad range of immunogenicity by conventional metrics. Median follow-up was 2.5 years after initial specimen collection. In total, 179 fresh tissue samples (74 tumor samples, 56 distant normal samples, 36 draining lymph nodes, and 13 metastasis samples) were collected from diagnostic laparoscopy (DL) with esophagogastroduodenoscopy (EGD) or gastrectomy for gastric adenocarcinoma by a specialist gastric cancer surgeon (**Methods**) (**Fig. 1a,b**). All samples were dissociated into single cell suspensions and viably cryopreserved within 2.5 hours of resection.

**Figure 1.**
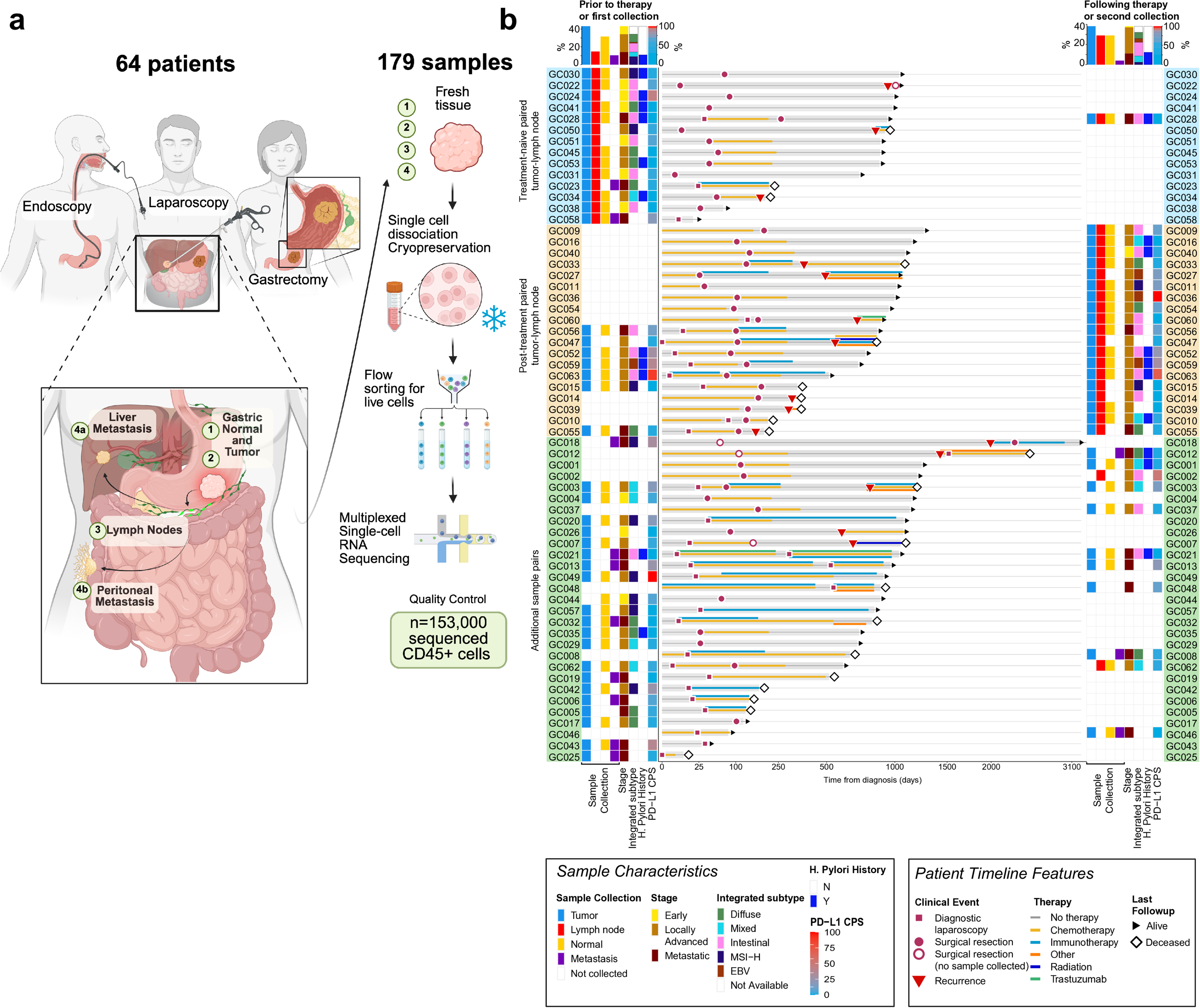
Patient cohort and sample collection. (a) Schematic of surgical sample collection from gastric cancer patients and processing for combined single cell RNA-seq, TCR-seq and surface epitope analysis. (b) Swimmer’s plot showing individual patients with samples collected, tumor characteristics, and simplified patient timelines. Samples are organized by those with paired tumor and dLN collections, and ‘additional sample pairs’ without both tumor and dLN.

### Distinct single cell immune profiles of gastric tumors and matched draining lymph nodes

We first analyzed immune cell subsets from 152,981 CD45+ cells passing quality control (QC) filtering (**Methods, Extended Data Table 3**). Unsupervised graph-based clustering identified six main CD45+ cell types marked by canonical identifying genes (**Fig. 2a, Extended Fig. 1a**): 98,274 T/NK cells, 23,560 B cells, 19,774 plasma cells, 8,876 myeloid cells, 1,687 mast cells, and 810 plasmacytoid dendritic cells (DCs). Further phenotypic subsets of CD8+ T cells (**Fig. 2b**), CD4+ T cells (**Fig. 2c**), and myeloid cells/DCs (**Fig. 2d**) were determined using a combination of known marker genes and surface proteins (**Extended Data Table 4**). Classical CD8+ T cell phenotypes including naïve, early activated, late activated, effector, central memory, proliferating, early exhausted, progenitor exhausted, terminally exhausted, effector memory, heat shock protein (HSP) associated and effector memory with expression of CD45RA (EMRA) subsets were identified, as well as CD8+ T cells marked by expression of *NEAT1* (**Extended Fig. 1b**). CD4+ T cell phenotypes included naïve, central memory, effector memory, follicular helper, cytotoxic, HSP-associated, proliferating, and Th17 cells, as well as a *NEAT1*-expressing cluster and two subsets of regulatory T cells distinguished by *CCR8* expression (**Extended Fig. 1c**). Proliferating CD4+ and CD8+ T cells were also identified (**Extended Fig. 1d**). Within the myeloid compartment, we identified mast cell and dendritic cell subsets including cDC1, cDC2, migratory DCs, and plasmacytoid DCs. We observed two macrophage populations, distinguished by either NLRP3+ inflammasome activation or expression of complement proteins (**Extended Fig. 1e**).

**Figure 2:**
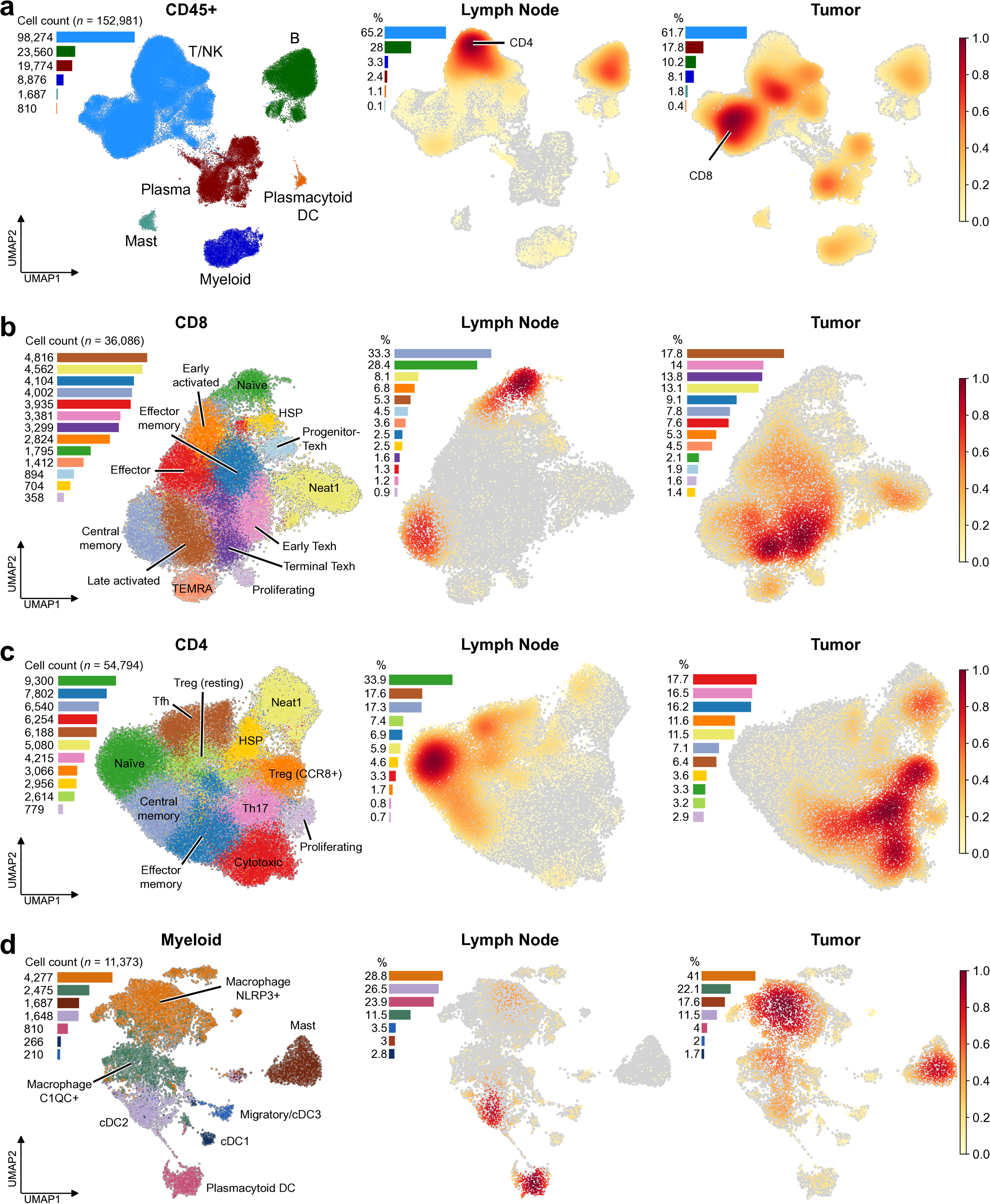
Single cell characterization of immune cells from primary tumor, distal normal, draining lymph nodes, and metastases in GC. Uniform manifold approximation and projections (UMAP) depicting (a) broad CD45+ cell subsets (left), density heatmap of CD45+ cells from dLN (center) and tumors (right), (b) CD8+ T cell subsets (left), density heatmap of CD8+ cells from dLN (center) and tumors (right), (c) CD4+ T cell subsets (left), density heatmap of CD4+ cells from dLN (center) and tumors (right), and (d) myeloid cell subsets (left), density heatmap of myeloid cells from dLN (center) and tumors (right).

Given the repertoire of immune cell phenotypes, we next quantified their relative distributions, comparing tumors to their matched dLNs. Lymph node and tumor samples were largely non-overlapping across all three cell compartments (CD8+, CD4+ and myeloid). CD8+ T cells were the more abundant immune cell type within tumors, whereas CD4+ T cells dominated both tumors and dLN (**Fig. 2a, Extended Fig. 1f**). Amongst CD8+ T cells, dLN primarily contained naïve and central memory cells while tumors contained primarily activated, effector memory, and exhausted T cells (**Fig. 2b**). CD4+ T cells within dLN were primarily naïve while tumors contained a mix of regulatory, Th17, and cytotoxic CD4 T cells (**Fig. 2c**). Finally, within the myeloid compartment, lymph nodes contained predominantly cDC2s and plasmacytoid DCs, while tumors were marked by NLRP3 macrophages and mast cells (**Fig. 2d**). Heat maps for the distribution of immune cells in normal and metastasis samples are in **Extended Fig. 2**. Thus, gastric tumors and draining lymph nodes comprise a non-overlapping immune network within which the distribution of cell types may enable or impair anti-tumor immune surveillance.

### Patient-specific immune networks defined by cytokine-driven cell-cell communication

Cytokines play an essential role in regulating the survival, trafficking, and function of immune cells across tissues^13^. Given the distinct immune cell composition of tumors and draining LN as well as the established role of chronic inflammation in gastric carcinogenesis, we set out to quantitatively measure cytokine-driven interactions between immune cells within gastric tumors and draining lymph nodes. Our goal was to phenotype LN and tumors based on their cytokine-immune crosstalk. Using a recently generated cytokine dictionary of gene expression changes in individual immune cell types isolated from mice treated with a library of 86 distinct cytokines^14^ (**Fig. 3a**, **Methods**), we quantified activation of gene expression programs associated with cytokine exposure in each cell type from our cohort. Briefly, after mapping cell types identified in **Fig. 2** to their murine counterparts, we computed cytokine enrichment in each cell type using an over-representation analysis at the sample level (**Methods**). We applied a newly developed Bayesian sparse tensor factorization approach (FRACTAL - Factor and Receiver Analysis of Cytokines in Tumor And Lymph nodes), effectively modeling variation in cytokine-driven expression programs across samples. This resulted in five robustly enumerated factors, with model weights in both ‘receiver’ cell-type and ‘cytokine’ activation dimensions (**Extended Fig. 3a**). Factor 0 was strongly associated with cDC2 activation, and moderately with pDCs. Factor 1 was strongly associated with Treg activation, and (γδ) T cells. Factor 2 was associated with CD4 T cells and Tregs, Factor 3 with macrophages, and Factor 4 with CD8+ T cells, NK cells, and B cells. Factors 0 and 2 were enriched in the dLN, while Factors 3 and 4 were enriched in the tumor; this is consistent with dendritic cell and CD4+ T cell abundance within lymph nodes, and macrophage and CD8+ T cell abundance within tumors. Regarding cytokines, IL-1a was enriched in Factor 2 while IL-15 and TNF were enriched in Factor 3. IL-1 alarmin was the highest activated cytokine family in Factors 1, 2, and 3, while Factors 0 and 4 were more balanced with the highest activity from the IL-2 family. Growth factors and receptor antagonists were the least activated cytokine families across all Factors.

**Figure 3.**
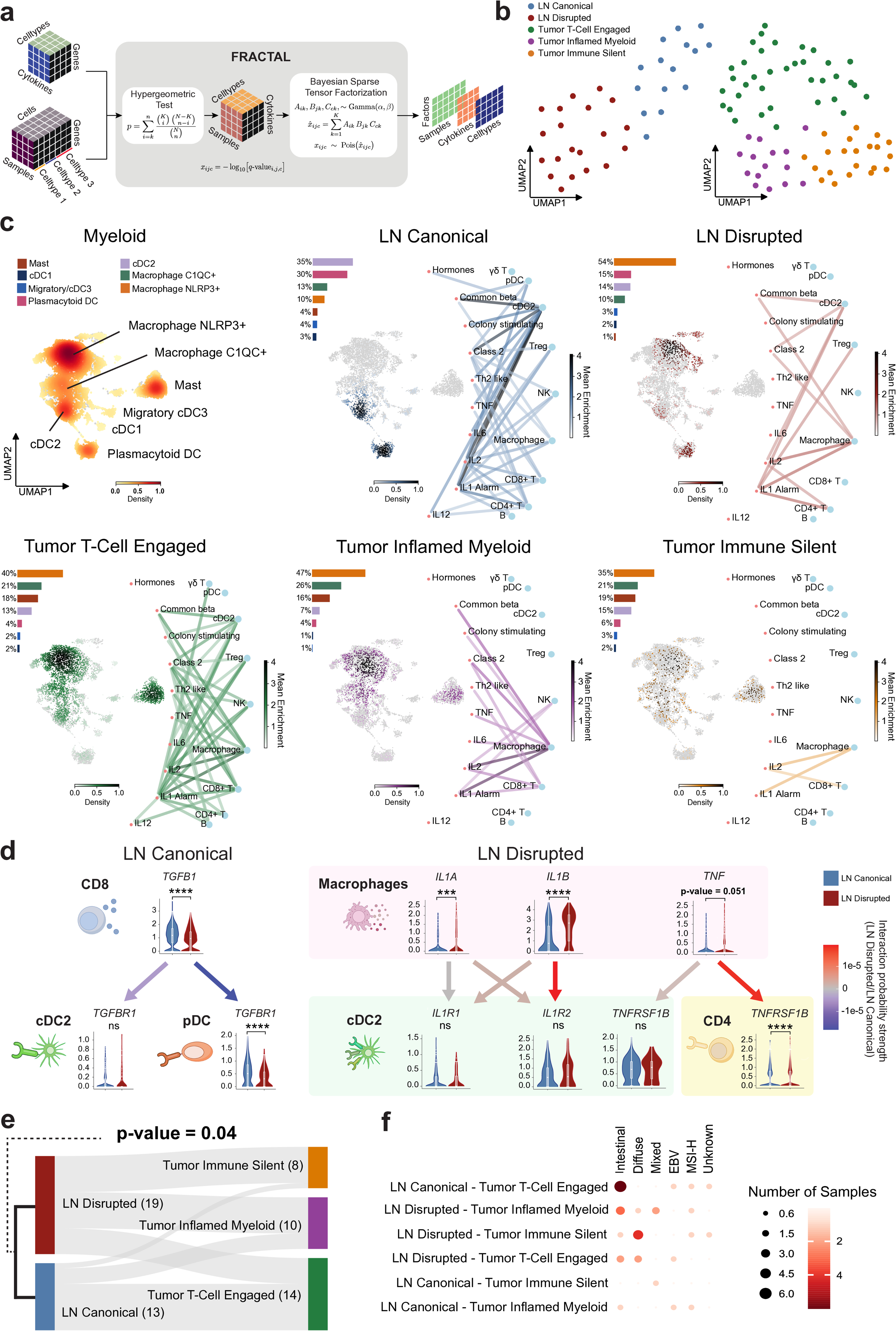
Patient stratification by distinct cytokine driven immune networks in tumors and their draining lymph nodes. (a) Workflow of FRACTAL - Factor and Receiver Analysis of Cytokines in Tumor And Lymph nodes our sample level Bayesian sparse tensor decomposition method. (b) Clustering of individual patient samples from dLN (left) and tumors (right) based on FRACTAL decompostion from (a). (c) Summary of enriched cytokine-cell type interactions in dLN and tumor subtypes with UMAPs showing the fraction of myeloid cell type content and paired myeloid cell density. (d) Significant cytokine ligand-receptor interactions identified using CellChat, with accompanying normalized gene expression. Interaction probability strength is shown relative to the disrupted lymph node samples, with blue arrows indicating interactions more probable in the canonical and red arrows more probable in the disrupted lymph node samples. (e) Correlation of cytokine interaction-dependent dLN and tumor subtypes in patients with paired samples demonstrates a significant association between dLN and tumor types as indicated (p-value = 0.04, chi-squared test) (f) Distribution of Lauren and molecular subtypes across paired dLN-Tumor networks.

We next stratified dLN and tumor samples by factor enrichment, yielding two clusters of cytokine-driven immune networks in dLN, and 3 clusters in tumors (**Methods**) (**Fig. 3b**). One dLN cluster (“canonical” LN) was defined by crosstalk between classical dendritic cells and T cells. Consistent with intact LN priming events, central memory CD8+ T cells, follicular helper CD4+ T cells, and type 2 and plasmacytoid dendritic cells were prevalent in this cluster (**Fig. 3c**, **Extended Fig. 3b**). The second dLN cluster (“disrupted” LN) showed a decrease in cytokine-driven transcriptional activity, with macrophages as the dominant cytokine-responsive cell type (**Fig. 3c**, **Extended Fig. 3b**). Notably this cluster harbored an abundance of CD8+ and CD4+ T cells with a naïve phenotype (**Extended Fig. 3b**) and macrophages with an NLRP3+ phenotype (**Fig. 3c**). Given the relative dominance of macrophage-driven crosstalk and the absence of dendritic cells needed for classical T cell priming, we labeled LN within this cluster as “disrupted”. Tumor clusters were similarly interpreted based on cytokine-driven immune cell interactions, with “T cell engaged” dominated by high-strength T cell interactions, activated and exhausted CD8+ T cells, and Th17 CD4+ T cells, “inflamed myeloid” with primarily NLRP3+ macrophage driven interactions, and “immune silent” with minimal immune cell infiltration or cytokine-driven interactions (**Fig. 3c, Extended Fig. 3b**).

We next asked which ligand-receptor interactions contributed most strongly to these cytokine-driven phenotypes. We inferred intercellular ligand-receptor (L-R) interactions, with a focus on cytokine-driven L-R interactions^15,16^ (**Methods**). Inference of interactions was consistent with dendritic cells as the dominant target of cytokine-driven L-R interactions within canonical LN, integrating signals from several immune cell subsets. Analogously, macrophages were the dominant source of cytokine-driven L-R interactions, broadly targeting immune cell subsets within disrupted LN (**Fig. 3d**). Transforming growth factor beta (*TGFB1*) was the dominant source of cytokine-mediated crosstalk within canonical LN, whereas several innate inflammatory cytokines, including IL-1 (*IL1A, IL1B*) and TNF, dominated cytokine-mediated crosstalk in disrupted LN samples (**Fig. 3d**).

We then assessed whether paired tumors and their dLN showed co-varying cytokine-driven reprogramming. The two dLN types led to significantly different tumor type distributions (p-value = 0.04, chi-squared test); canonical dLN were more often paired with T-cell engaged tumors, while disrupted dLN were more often paired with either immune silent or inflamed myeloid tumors (**Fig. 3e**). Individual patient tumor and dLN types are listed in **Extended Data Table 5**. There was no clear correlation of tumor genomic features with either tumor or dLN type (**Extended Fig. 3c**). Interestingly, there was an uneven distribution amongst traditional tumor types, with diffuse-type tumors, which carry the poorest prognosis amongst gastric tumor subtypes^17^, pairing exclusively with disrupted dLN (**Fig. 3f**). As T cell priming occurs in the lymph node, these results suggest that altered priming within dLN contributes to the lack of intratumoral T cell surveillance in these tumor subtypes.

### Reduced T cell priming in disrupted tumor draining lymph nodes

Linked cytokine-driven reprogramming between tumor and matched dLN suggests that chronic inflammation-driven loss of dendritic cell homing to disrupted dLN might impair adaptive immune responses. To investigate this, we next focused on the impact of canonical and disrupted dLN on CD8+ T cell phenotypic trajectories. We generated a CD8+ T cell trajectory manifold (**Methods**) and mapped T cells according to their transcriptional phenotypes (**Fig. 4a**). T cells from dLN and tumors had limited phenotypic overlap (**Fig. 4b**). A computational fate analysis (**Methods**) revealed 6 primary terminal states for naïve CD8+ T cells, including activated, effector, EMRA, Neat1+, and proliferating fates, as well as a cell fate in which T cells remained in a naïve-like state (**Fig. 4c**). Using single-cell TCR analysis, we confirmed that clonally expanded T cells were mostly, though not exclusively, in the tumor (**Extended Fig. 4a**). The tumor-resident clonally expanded T cells were comprised of a wide phenotypic range of effector and exhausted states, while dLN expanded T cells were primarily in a central memory state (**Extended Fig. 4a**).

**Figure 4.**
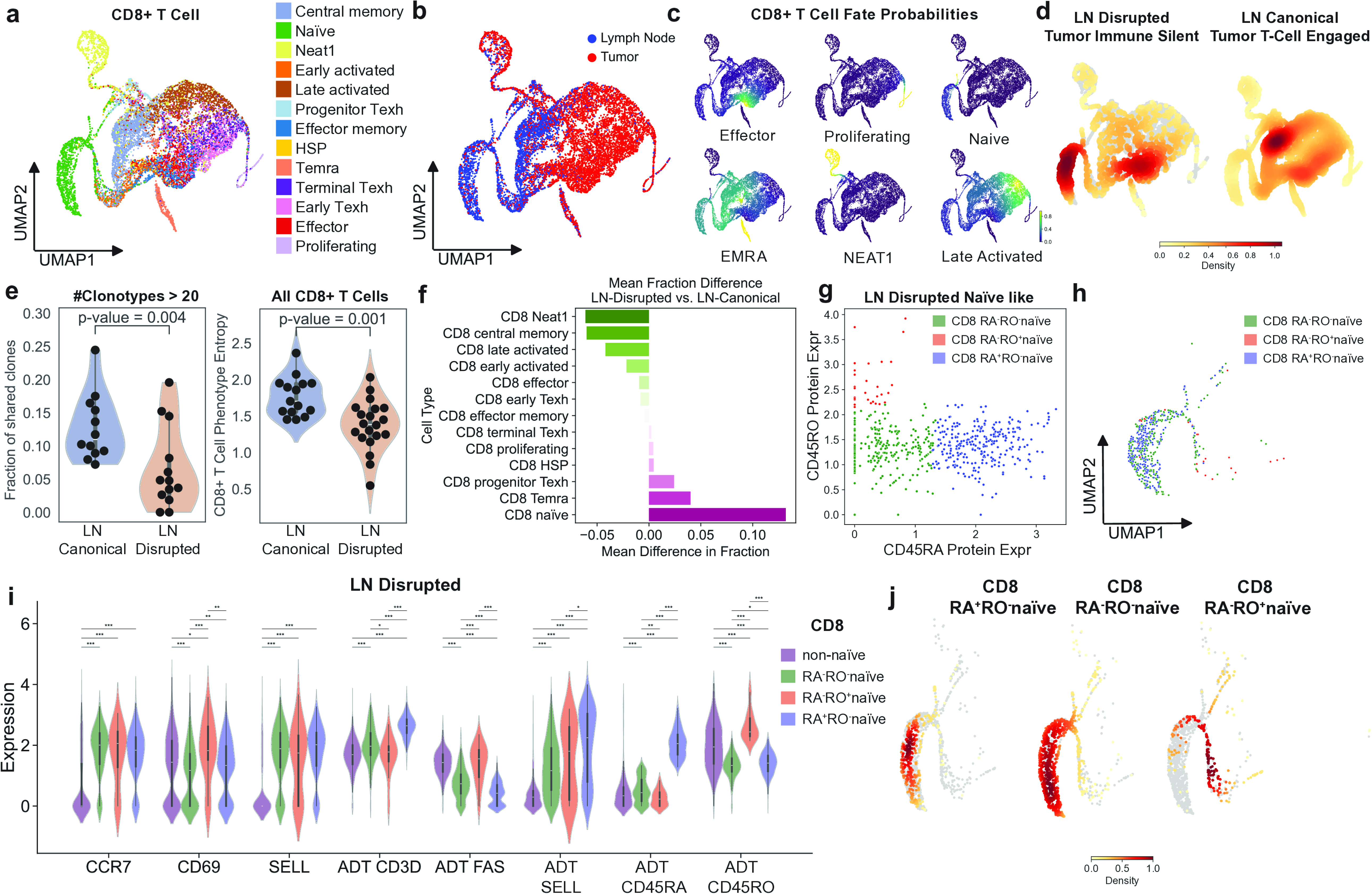
Reduced T cell priming in disrupted tumor draining lymph nodes. (a-d) Trajectory manifold of CD8+ T cell subsets across dLN and tumor, showing distribution of (a) CD8+ T cell subsets, (b) distribution within dLN and tumors, (c) fate probabilities, and (d) density mapping of cells from ‘disrupted-immune silent’ pairs and ‘canonical-T cell engaged’ pairs. (e) Fraction of shared expanded (clonotypes >20) CD8+ T cell clones between dLN and Tumor (left) and average phenotype entropy in CD8+ T cells (right) (**Methods**, section *Phenotype entropy*), stratified by canonical dLN versus disrupted dLN. (f) Difference in mean cell fractions of CD8+ T cell subsets between disrupted dLN and canonical dLN. (g) Surface CD45 isoform expression distribution on CD8+ T cells from disrupted dLN with a transcriptionally naïve phenotype. (h) Cells from (g) projected on the trajectory manifold from (a), coded by CD45 isoform expression. (i) Expression of genes and proteins associated with T cell activation on phenotypically naïve T cells from (g,h), stratified by surface expression of CD45 isoforms. (j) Density heatmaps of subtypes of naïve cells from (g-i), projected onto the CD8+ trajectory manifold from (a).

We next focused on comparing T cell differentiation trajectories from canonical and disrupted LN priming. In canonical LN-tumor pairs, dLN CD8+ T cells were primarily comprised of a central memory phenotype and were found in all six terminal states within the tumor. In sharp contrast, T cells from disrupted LN-tumor pairs were restricted to undifferentiated naïve-like states within the dLN, and their tumor resident trajectories were substantially more restricted (**Fig. 4d**). We attributed this constrained differentiation directly to altered priming within the dLN, as TCR analysis revealed that CD8+ T cells within disrupted LN displayed reduced clonal expansion and a reduced phenotypic entropy (**Fig. 4e**), with the majority of CD8+ T cells occupying either a naïve or EMRA phenotypic state (**Fig. 4f**).

Curiously, a fraction of phenotypically naïve CD8+ T cells from disrupted dLN downregulated surface expression of CD45RA, which typically occurs in the context of antigen experience^18,19^ (**Fig. 4g**), despite being transcriptionally similar to naïve cells (**Fig. 4h**). We therefore stratified phenotypically naïve T cells based on CD45 isoform expression (CD45RA vs. CD45RO, included in our CITE-seq antibody-derived tags (ADT) panel). Phenotypically naïve T cells negative for CD45RA and positive for CD45RO (‘RA^-^RO^+^’) expressed markers consistent with antigen experience on the protein level, including downregulation of L-selectin and CD3^20,21^, as well as upregulation of Fas^22^, consistent with being recently activated. In contrast, T cells negative for CD45RA and CD45RO (‘RA^-^RO^-^’) largely overlapped with CD45RA+, CD45RO-canonically naïve T cells, with sustained expression of L-selectin and CD3 (**Fig. 4i**). Trajectory analysis of these 3 phenotypically naïve T cell subsets along with early activated T cells confirmed that RA^-^RO^-^ T cells were similar to RA^+^RO^-^ naïve T cells, whereas RA^-^RO^+^ cells were more similar to early activated T cells (**Fig 4j, Extended Fig. 4b**). We therefore combined double negative RA^-^RO^-^ T cells with RA^+^RO^-^ true naïve cells and RA^-^RO^+^ cells with early activated T cells for downstream analysis. We found that compared to canonical dLN, T cells within disrupted dLN were enriched in either a naïve or EMRA phenotype, indicating that inflammatory reprogramming of disrupted dLN is sufficient to limit T cell priming and downstream tumor surveillance (**Extended Fig. 4c**).

### KLF2 suppresses effector T cell differentiation in gastric tumor-draining LN

Despite showing increased clonal expansion relative to T cells from disrupted LN, T cells from canonical LN also showed evidence of phenotypic constraint, with the majority of clonally expanded T cells within dLN remaining in a central memory state (**Extended Fig. 5a**). We therefore sought to identify transcriptional regulators that might restrict differentiation and egress of CD8+ T cells from dLN. We identified differentially expressed transcription factors in clonally expanded CD8+ T cells compared with clonally expanded T cells from tumors and found Kruppel-like Factor-2 (*KLF2*)^23^ as the most differentially upregulated transcription factor in clonally expanded T cells retained within dLN (**Fig. 5a**). Notably, within both dLN and tumors, *KLF2* expression correlated strongly with expression of genes linked to quiescence (*LEF1, TCF7*) and LN retention (*S1PR1, S1PR4, CCR7, SELL*) and inversely with genes linked to tissue infiltration (*ITGAE*, *CXCR6*) and effector function (*IRF4, GZMB*) (**Fig 5b**). Moreover, as *KLF2* expression inversely correlated with clone size amongst expanded clones within the LN and tumor, we inferred that *KLF2* expression may be a regulator of quiescence and LN retention while restricting T cell expansion, differentiation, and LN egress (**Fig. 5c**).

**Figure 5.**
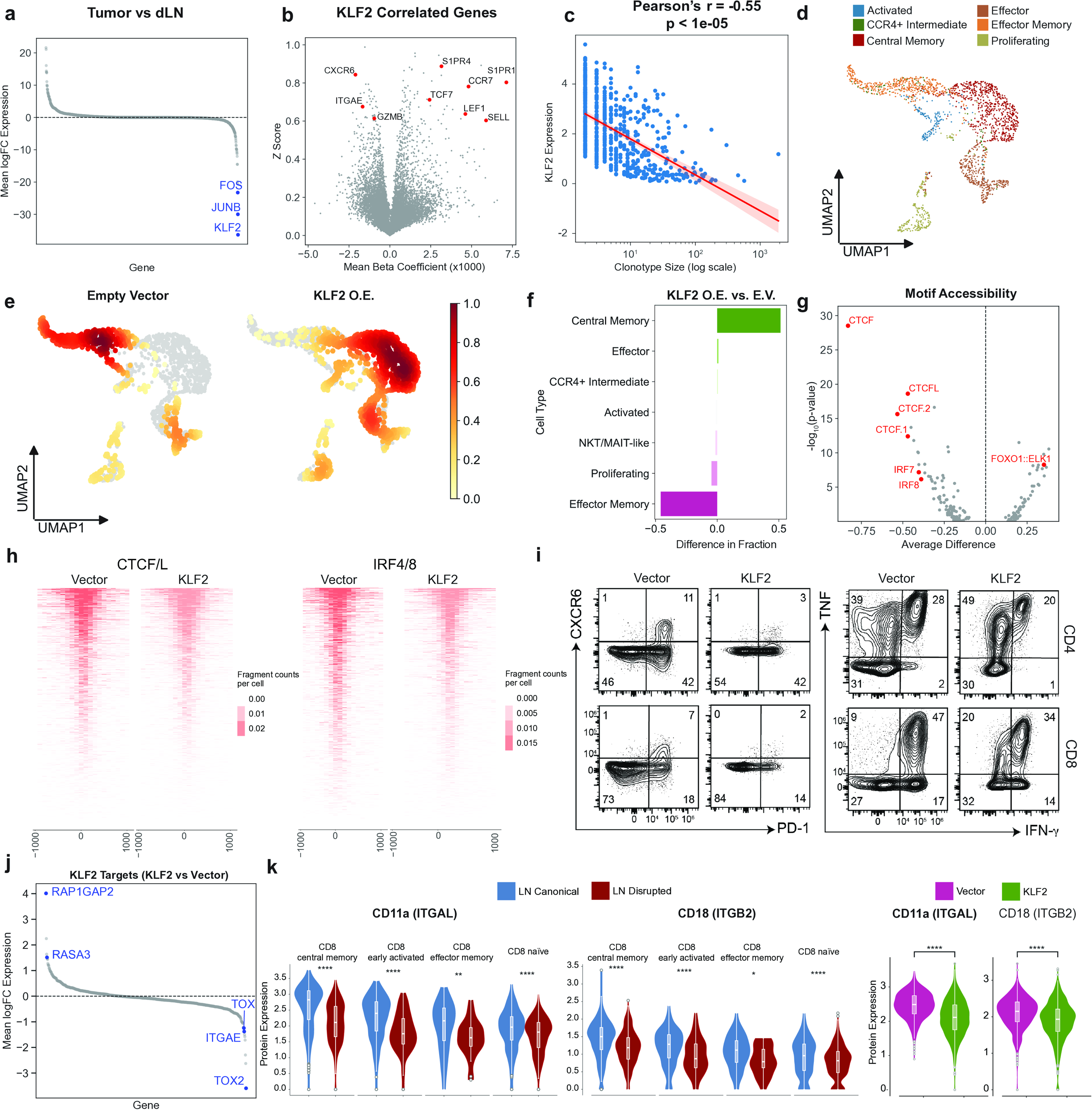
KLF2 suppresses effector T cell differentiation. (a) Differentially expressed transcription factors expressed in non-naïve CD8+ T cells from tumors (up) vs. dLN (down). (b) Genes correlated with the KLF2 expression in non-naïve CD8+ T cells across tumors and dLN (c) Correlation of KLF2 expression on non-naïve CD8+ T cells across tumors and dLN with clonotype size. (d) UMAP of activated human CD8+ T cells transduced with either empty vector or a KLF2 overexpression vector. (e) Density plot showing relative distribution of CD8+ T cell phenotypes in primary human CD8+ T cells transduced with either empty vector (left) or KLF2 overexpressing vector (right). (f) Mean fraction of T cell subsets in primary human CD8+ T cells transduced with either empty vector or KLF2 overexpressing vector as indicated. (g) Transcription factor motifs with top accessibility scores in central memory (right) compared with effector memory (left) T cell subsets. (h) Volcano plot demonstrating accessibility of CTCF/L (left) or IRF4/8 (right) motifs, centered around the transcriptional start site, in human CD8+ T cells transduced with empty vector or KLF2 overexpression vector. (i) Expression of CXCR6 and PD-1 (left) and intracellular accumulation of IFN-g and TNF following restimulation with PMA and ionomycin in CD4+ and CD8+ T cells transduced with empty vector or KLF2 overexpression vector. *ADT* antibody-derived tags (j) Relative expression of genes (KLF2 overexpression vector vs. Empty vector) with an overlapping KLF2 motif (k) Differential surface expression of the LFA-1 subunits CD11a and CD18 in indicated CD8+ T cell subsets from dLN, stratified by canonical vs. disrupted dLN phenotype (left) and in primary human T cells transduced with empty vector or KLF2 overexpression vector.

To directly assess the consequences of *KLF2* stabilization on T cell activation and differentiation, we transduced activated primary human T cells with either lentivirus encoding *KLF2* or a vector control (**Extended Fig. 5b**) and determine the impact of *KLF2* overexpression on T cell phenotypes using a combination of flow cytometry and combined single-cell RNA and ATAC sequencing (DOGMA-Seq)^24^. *In vitro* activated T cells differentiated into several T cell states marked by distinct transcriptional modules, including proliferating, effector, effector memory, and central memory T cells, with a minority of T cells occupying innate-like, activated, or intermediate states (**Fig 5d, Extended Fig. 5c**). Naïve cells were not present as T cells had been previously activated to facilitate lentiviral transduction. Consistent with *KLF2* expression and accessibility marking a central memory phenotype^25^, we found that overexpression of *KLF2* in primary T cells led to an increased abundance of CD8+ T cells with a central memory phenotype and a decreased abundance of T cells with an effector memory phenotype (**Fig. 5e,f**). To determine the mechanism by which *KLF2* alters T cell differentiation, we identified differentially accessible motifs in central memory versus effector memory T cells using ChromVar^26^. We found that CCCTC-binding factor (*CTCF*) and Interferon Regulatory Factor (*IRF*) motif accessibility were associated with effector memory T cells, whereas *FOXO1* and T-box (*TBX*) family members were associated with central memory T cells (**Fig. 5g**). Measurement of promoter-centered chromatin accessibility confirmed that promoter-associated accessibility of both *CTCF* and *IRF* transcription factors was reduced in *KLF2*-overexpressing cells (**Fig. 5h**). As both *CTCF* and *IRF* transcription factors are known to be required for effector CD8+ T cell differentiation^27–32^, we measured both effector T cell differentiation and cytotoxic function and found them to be reduced in *KLF2* overexpressing T cells (**Fig 5i, Extended Fig. 5d**).

Finally, to identify potential mechanisms by which *KLF2* might limit T cell differentiation and tumor infiltration, we identified putative *KLF2* target genes by asking which genes containing a KLF-binding motif^33^ (CCACGCCC) were differentially expressed in *KLF2* versus vector overexpressing CD8+ T cells. *KLF2* overexpression was associated with reduced expression of target genes associated with effector memory T cells, such as *TOX*, *TOX2*, and *ITGAE* and increased expression of the GTPase activating proteins (GAP) *RAPGAP2* and *RASA3* (**Fig. 5j**). *RAPGAP2* and *RASA3* bind and stimulate the catalytic activity of the Rap1 GTPase^34–36^. Rap1 has been shown to be required for the TCR-dependent conformational change in Lymphocyte function-associated antigen-1 (LFA-1) that enables interaction with its ligands, otherwise known as ‘inside out’ signaling. This led us to hypothesize that *KLF2*-driven expression of Rap1 GAPs may limit T cell priming by disrupting LFA-1 activation. Consistent with this hypothesis, surface expression of both components of the LFA-1 heterodimer (CD11a and CD18) was reduced in naïve and central memory T cells from disrupted LN; moreover, *KLF2* expression *in vitro* was sufficient to lower expression levels of CD11a and CD18 when compared to our empty vector control (**Fig. 5k**). To directly test the impact of LFA-1 activity on T cell priming and differentiation, we activated OT-I TCR transgenic T cells with cognate peptide across a range of antigen dose and affinities using altered peptide ligands of the canonical OT-I TCR cognate peptide SIINFEKL (**Extended Fig. 6a**)^37^. Addition of LFA-1 blocking antibodies^38^ during T cell priming by peptide-pulsed DCs increased the threshold for activation, proliferation, and effector function across peptide doses and affinities, providing a mechanism by which *KLF2* stabilization influences both T cell priming and effector differentiation in GC (**Extended Fig. 6b,c**).

### Dynamic state switching of cytokine driven dLN types in a remarkable patient

Lymph nodes are typically only surgically resected during gastrectomy, as a fundamental part of staging. However, for one specific patient (GC028) a biopsy of a visible draining lymph node was collected during diagnostic laparoscopy (DL) for evaluation of metastatic gastric cancer as well as during surgical resection after achieving a near complete response to neoadjuvant chemotherapy (FOLFOX)^39^ (**Fig. 6a**). This unique pairing of tumor and draining lymph node both before and after neoadjuvant chemotherapy allowed us to investigate whether cytokine driven dLN types remain fixed in the context of therapy. Classification of both pre-treatment and post-treatment samples using FRACTAL revealed a disrupted dLN prior to treatment and a canonical dLN at the time of resection (**Fig. 6b**). Accordingly, the post-treatment dLN showed a decreased abundance of C1QC+ macrophages and an increased abundance of plasmacytoid DCs (**Fig. 6c, Extended Fig. 7a**).

**Figure 6.**
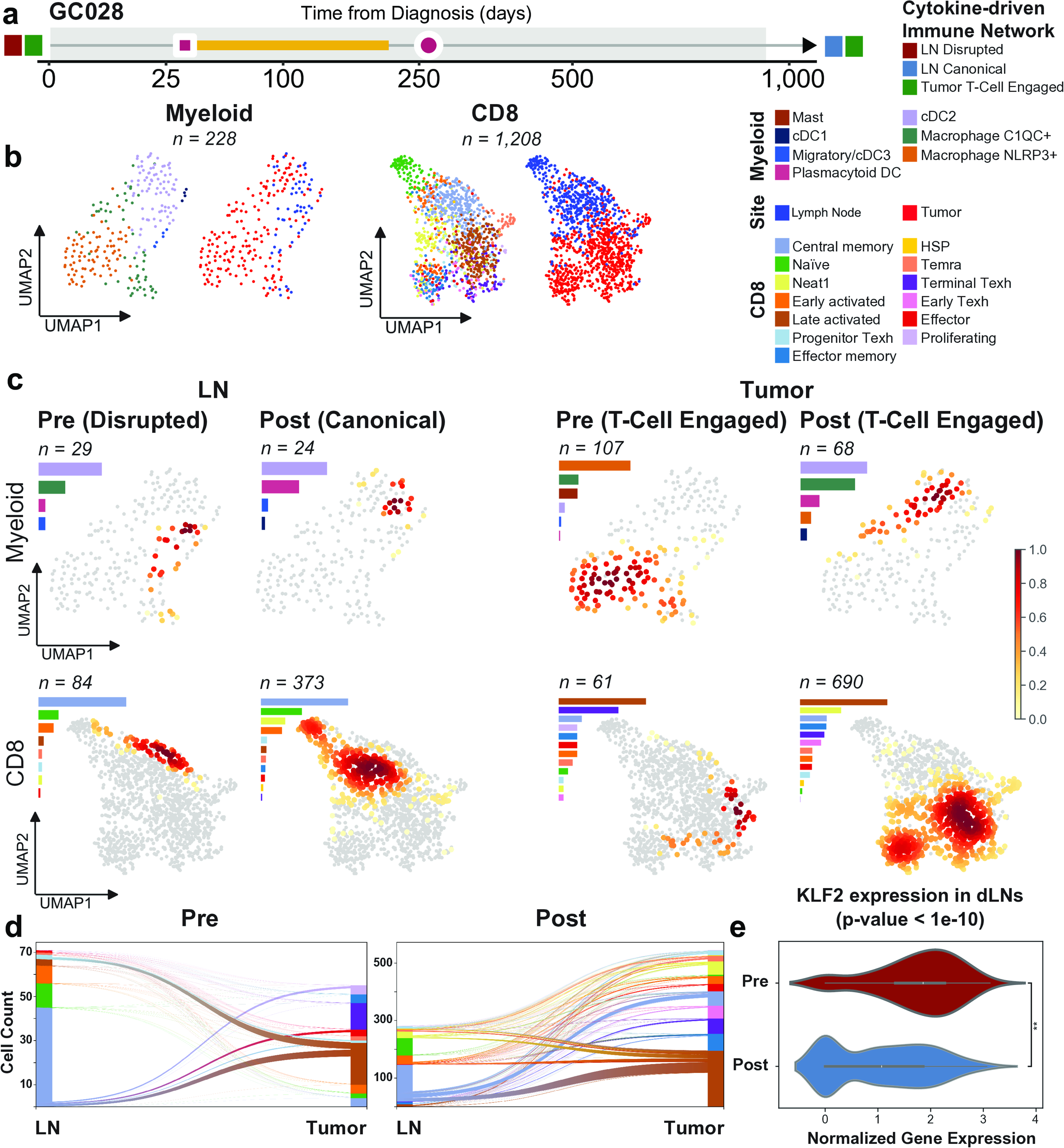
Cytokine driven dLN types are not fixed. (a) Patient timeline with tumor and LN collection pre- and post-neoadjuvant chemotherapy treatment. (b) UMAP projections of myeloid cells (left) and CD8+ T cells (right) for patient GC028 across both pre-treatment and post-treatment conditions, colored by either cell subset or distribution in tumors and dLN. (c) Heat-encoded density embeddings of myeloid and CD8+ T cell compartments prior to and following neoadjuvant treatment, stratified by timepoint and anatomical site. (d) Alluvial plots showing CD8+ T cells with shared TCRs across dLN and tumor, both pre- and post-treatment (e) KLF2 expression in CD8+ T cells within dLN of patient GC028, pre- and post-treatment.

Clonotype tracking between the dLN and tumor pairs revealed that shared clones were present between all 4 samples (**Extended Fig. 7b**). Phenotypes of shared CD8+ T cell TCR clonotypes changed post-treatment, with an increase in the fraction of progenitor Texh in the tumor and CM in the dLN (**Extended Fig. 7c**). The post-treatment canonical dLN also had both a larger number of expanded clones and a greater diversity of expanded T cell phenotypes from the dLN to its paired tumor (**Fig. 6d**). We examined overall *KLF2* expression in CD8+ T cells in the pre-therapy disrupted dLN and the post-therapy canonical dLN and found that *KLF2* expression was significantly higher in the disrupted dLN (pre) than the canonical dLN (post) (**Fig. 6e**). *KLF2* was also the second most transcriptionally decreased gene between these pre-treatment and post-treatment dLN (**Extended Fig. 7d**). Notably, expression of both LFA-1 components (CD11a and CD18) was also increased in the canonical dLN (post-treatment) (**Extended Fig. 7e**). These observations imply that the disrupted and canonical types we report above are not fixed in disease evolution. Rather, changes in *KLF2* expression coupled with cytokine driven dLN dynamics suggest therapeutic intervention can modulate lymph node priming in gastric cancer. The clinical and prognostic significance of this single patient remains unclear, however we note that this patient remains in complete remission 2.7 years after diagnosis despite presenting with metastatic disease.

## Discussion

Our study sought to understand the impact of chronic inflammation on lymph node priming events during gastric tumorigenesis. Through multi-modal single cell investigation of gastric tumors and matched draining lymph nodes, we provide here a novel mechanistic explanation for how persistent innate inflammation can impair anti-tumor T cell immunity. Our findings are consistent with reduced dendritic cell accumulation within disrupted draining lymph nodes, preventing exposure of putatively tumor-reactive naïve T cells within the lymph node to tumor-derived neoantigens. While diffuse-type GC had exclusively disrupted dLN, we identified this LN phenotype across every molecular GC subtype, indicating that LN disruption suppresses immune surveillance in GC across histologic and molecular classifications. These findings nominate DC homing and T cell priming as potential therapeutic targets that cross previously reported GC stratifications.

We also identify a second, previously unappreciated checkpoint on T cell surveillance in gastric cancer. Activated T cells within the draining LN largely fail to differentiate into effectors, instead becoming central memory T cells that remain in the LN and express high levels of *KLF2*. This may be in part due to reduced integrin activity increasing the threshold for T cell activation during naïve T cell priming events within dLN. These findings are supported by recent evidence demonstrating a role for *KLF2* in preventing T cell exhaustion^40^. By suppressing the exhaustion program*, KLF2* may paradoxically reduce the clinical efficacy of immune checkpoint inhibitors that target inhibitory receptors preferentially expressed on exhausted T cells, including PD-1, CTLA-4, and LAG-3. Future studies to determine if *KLF2* expression in T cells is associated with lack of immune checkpoint blockade (ICB) response or whether targeting either *KLF2* or *KLF2*-dependent integrin activation can enhance sensitivity to ICB would have immediate clinical relevance.

Our findings provide insights into how reciprocal balances between innate and adaptive immune responses maintain homeostasis in an inflammatory microenvironment, and how an imbalance between these responses contributes to human disease. The extent to which the T cell phenotypes observed in this study apply to other tumor types remains unknown. However, this unique dataset, derived from prospective surgical sampling of paired tumors and draining LN, may serve as a valuable reference to understand whether lymph node priming is disrupted in other cancers. Blunting of the adaptive immune response during chronic inflammation may represent a physiologic mechanism to preserve mucosal tolerance and barrier integrity; consistent with this hypothesis, T cell specific *KLF2* expression is required to maintain commensal tolerance and prevent inflammatory bowel disease^41^. However, adaptive immune suppression has been reported in other chronic inflammatory conditions, including sepsis-associated immune paralysis^42^. Potential targets such as *KLF2* may enable T cells to escape from this maladaptive state. Furthermore, our anecdotal evidence that dLN can change with intervention supports tumor draining lymph nodes as a viable focus for therapeutic development. We report that a patient with pre- and post-treatment tissue collections demonstrated phenotypic T cell state transitions linked to a remarkable clinical response. This indicates that the non-canonical immune states we describe in this study are potentially reversible with therapeutic intervention. Characterizing the prevalence and determinants of such immune state transitions will be central to future translational efforts aimed at enhancing lymph node priming in gastric cancer.

## Extended Figure Legends

**Extended Figure 1. Marker genes for major CD45+ cell compartments.** The expression of marker genes in each major CD45+ cell compartment (a), CD8+ T cells (b), CD4+ T cells (c), cycling cells in CD4+ and CD8+ T cells (d) and myeloid cells (e). (f) Fraction of cells in the major CD45+ cell compartments within tumors and dLN as indicated.

**Extended Figure 2. Single cell characterization of immune cells from normal tissue and metastatic sites in GC.** UMAPs depicting (a) broad CD45+ cell subsets across tissue sites (left), density heatmap of CD45+ cells from distant normal tissue (center) and metastases (right), (b) CD8+ T cell subsets across tissue sites (left), density heatmap of CD8+ cells from distant normal tissue (center) and metastases (right), (c) CD4+ T cell subsets across tissues (left), density heatmap of CD4+ cells from distant normal tissue (center) and metastases (right), and (d) myeloid cell subsets across tissue sites (left), density heatmap of myeloid cells from distant normal tissue (center) and tumors (right).

**Extended Figure 3. Patient stratification by distinct cytokine driven immune networks in tumors and their draining lymph nodes.** (a) Posterior mean for latent sample by factor (left), cell-type by factor (center), and cytokine by factor (right) tensor slices. For visualization purposes, the cytokine by factor slide heatmap includes only cytokines with an average factor activation score of more than 0.2. (b) UMAPs showing the fraction of CD8+ T cell type content and paired CD8+ T cell density (left) and fraction of CD4 and myeloid cell subsets in each cytokine interaction-driven immune network cluster. (c) Oncoplot of tumor mutation^43^, separated by either dLN or tumor subtype.

**Extended Figure 4. CD8+ T cell expansion.** (a) CD8+ T cell clonotype size distribution in dLN and Tumor (left) and cell phenotypes for expanded TCR clonotypes (right). (b) Average distance of naïve like T cells from non-naïve T cell phenotypes indicating closer proximity of RA-RO+ naïve T cells to early activated T cells (**Methods**). (c) Difference in mean cell fractions of CD8+ T cell subsets between disrupted dLN and canonical dLN following recategorization of naïve-like CD8+ T cells (RA^+^RO^-^ naïve and RA^-^RO^-^ naïve into ‘naïve merged’, RA^-^RO^+^ naïve and early activated into ‘early activated merged’).

**Extended Figure 5. KLF2 suppresses effector T cell differentiation.** (a) Fraction of CD8+ T cell phenotypes in expanded TCR clonotypes in each cytokine-driven immune network cluster. (b) Western blot showing KLF2 and Histone H3 (loading control) protein abundance in primary activated human T cells transduced with either empty vector control or KLF2 overexpression vector. (c) Marker genes and their expression in CD8+ T cell phenotypes reported in empty vector and KLF2 overexpression vector-transduced cells from Fig. 5d. (d) Flow cytometry gating for Fig. 5i.

**Extended Figure 6. LFA-1 blockade attenuates T cell priming and effector differentiation.** (a) Experimental design to assess the contribution of LFA-1 to DC-mediated T cell priming and effector differentiation. (b) Flow cytometry scatter plots and gating of live, B220^-^NK1.1^-^ TCRb^+^CD8^+^ T cells to assess T cell activation (CD69^+^CD25^+^), proliferation (CD44^+^CellTrace Violet^low^), effector differentiation (CX3CR1^-^CXCR6^+^) and cytokine production (TNF^+^IFN-g^+^). (c) Quantitation of impact of peptide dose and affinity on CD8+ T cell activation, proliferation, effector differentiation, and cytokine production as described in (b). Each **table entry** represents the **average** of three independent biological replicates for each peptide affinity and dose. *p<0.05 **p<0.01 ***p<0.001

**Extended Figure 7. Cytokine driven dLN types are not fixed** (a) Fraction of CD8+ T (left) and myeloid cell type subsets (right) in patient GC0028, stratified by tissue (dLN and tumor) as well as timepoint (pre- and post-treatment) (b) Abundance of TCR clonotypes that are shared across time and anatomical sites (left) and their projection onto the patient GC0028-specific CD8+ T cell UMAP (right) (c) Alluvial plots showing phenotypes of CD8+ T cells with shared TCR pre- and post-treatment in both dLN (left) and tumor (right) samples. (d) Differentially expressed genes in the dLN pre-treatment (up) vs post-treatment (down). (e) Expression of CD11a and CD18 in CD8+ T cells from patient GC0028 dLN pre- and post-treatment.

## EXTENDED DATA TABLES

**Extended Data Table 1**: Individual patient and tumor characteristics

**Extended Data Table 2**: Patient demographics

**Extended Data Table 3**: Data subsets and cell counts

**Extended Data Table 4**: Top 50 DEGs for cell type cluster annotation for CD45+, CD8, CD4 and Myeloid subsets

**Extended Data Table 5:** Patients with dLN and tumor IREA types

## END NOTES

## Acknowledgements

This research was supported in part by the institution’s Cancer Center Support Grant (P30 CA008748) from the NIH, and the Cycle for Survival-Equinox Grant (MA, EES, VES, SAV). SAV was supported by a Damon Runyon Clinical Investigator Award. SS is supported by an NCI Pathway to Independence Award (5 K99 CA277562-02). Additional support was provided by the Halvorsen Center for Computational Oncology. SPS holds the Nicholls-Biondi Endowed Chair in Computational Oncology. This work was additionally supported by research funding from Bristol Myers Squibb. Figure 1a and Extended Figure 6a were made using Biorender.com. We thank the MSKCC Single Cell Analysis and Innovation Lab (SAIL) for their thoughtful insights and assistance, Anthony Verdone for assistance with data management, and Monique Du Plessis for assistance with Figure preparation. This work uses resources from the High-Performance Computing Group at Memorial Sloan Kettering Cancer Center.

## Author Contributions

EES, SS, SS, MA, HL, VES, SPS, and SAV conceived the project and designed all experiments. SS, MA, HD, VES and EES performed all sample acquisition. EES and SS assembled all metadata and reviewed clinical characteristics. YHL, SS and EES performed all tissue processing. YHL and EES performed all spectral cytometry assays. YHL and EES performed all laboratory experiments. SS, HL, MZ, and NC performed all computational and statistical analyses. EES, SS, HL, SPS, VES and SAV wrote the manuscript; all authors reviewed the manuscript.

## Data and Code Availability

Supplementary Information is available for this paper. Scripts used in the analysis, including the implementation for FRACTAL is available online at https://github.mskcc.org/shahcompbio/gac_code.

## METHODS

### Patient Inclusion

Under Institutional Review Board approval (IRB 20-025) fresh tissue was collected from gastric tumors, distant normal gastric mucosa, draining gastric lymph nodes (dLN), and metastases from patients with gastric adenocarcinoma at Memorial Sloan Kettering Cancer Center (MSK). Patients had either institutional 6-107 and/or 12-245 tissue consent or specific IRB approval for tissue use after patient death. IRB approval was obtained for patients under 18. Tissues were collected from patients undergoing either gastric resection or diagnostic laparoscopy (DL) with esophagogastroduodenoscopy (EGD) as part of treatment or staging/diagnosis. Gastric resections were either open or robotic assisted. Biopsies taken during EGD were removed using cold endoscopic forceps, and biopsies taken during DL were removed using cold laparoscopic graspers. Pathology was reviewed by institutional gastrointestinal pathologists and all tumors were pathologically confirmed to be gastric adenocarcinoma. The 8^th^ edition of the AJCC staging system was used by convention for pathologic TNM staging.^1^ Clinical information for patients was collected prospectively in a clinical gastric database. Lauren classification^2^ and TCGA classifications^3^ were combined to describe patients with intestinal, diffuse, mixed, MSI-H and EBV tumors. MSI-H or EBV designation superseded Lauren classification. Lauren classification was determined by pathology report. MSI-H status was determined by immunohistochemistry (IHC) of MLH1, MSH2, MSH6, and PMS2 as reported by pathology or by tumor mutation burden on institutional genomic testing (MSK-IMPACT).^4^ EBV positivity was determined by IHC on pathology report. HER2 status was reported as negative, positive, or NA (not available) as a summary of HER2 IHC and fluorescence in situ hybridization (FISH) analysis for equivocal IHC results from pathology reports. PD-L1 scores were reported as CPS from 0-100, with any scores over 100 reduced to 100. For patients who received systemic therapy prior to resection, tumor response grade (TRG) was reported as called in the institutional pathology report from 0-3 according to the NCCN system.^5^ *H. pylori* history was determined by a previous positive IHC test or a documentation of previous *H. pylori* infection or treatment in a patient’s chart. Treatments were grouped as chemotherapy, immunotherapy, trastuzumab, radiation, or other and were categorized as prior to or following specimen collection. Survival, follow-up, and recurrence were collected up to January 17, 2025. Outcomes for two patients were not included due to their enrollment in a clinical trial after specimen collection. Disaggregated data could not be publicly shared for **7** patients due to IRB restrictions.

### Sample Collection and Processing

As mentioned above, tissue was collected in the operating room from clinically indicated gastric cancer resections or DLs with EGDs. During robotic gastric resections, Indocyanin Green (ICG) was injected intra-operatively to mark the tumor bed and highlight tumor-draining lymph nodes as previously described.^6^ During open operations, tumor draining lymph nodes were determined by the operating surgeon according to tumor location and lymph node station. For gastrectomies, within 3 minutes of resection, 0.5-1.0 cm^3^ of tumor, distant normal gastric mucosa (defined as macroscopically normal appearing to the surgeon and not directly adjacent to the tumor), and LNs (identified to be draining nodes by ICG or station by operating surgeon) were isolated, cut into ∼1 mm^3^ pieces and placed into 15mL conical tubes containing phosphate buffered saline (PBS). These samples were placed on ice and immediately transported to the laboratory. During DL with EGD tumor and distant normal samples were collected during EGD with endoscopic cold forceps, 4 bites of tumor were collected then immediately suspended in PBS, as were 3 bites of distant normal gastric mucosa with the forceps cleaned in-between collections. If peritoneal or liver metastases were seen on DL with sufficient tissue for pathologic diagnosis, a biopsy of 0.5 cm^3^ of metastasis was collected and immediately transferred to a 15 mL conical tube with PBS. These samples were placed on ice and immediately transported to the laboratory in the same manner as for gastric resections. All samples were digested using a GentleMACS Dissociator in 2.2 ml or 4.7 mL (based on tissue size) of R-10 solution (RPMI, 10% fetal bovine serum, 10mM HEPES, 1% penicillin-streptomycin, 1% L-glutamine) and Miltenyi enzymes (Miltenyi Biotec Tumor Dissociation Kit, Human) at 37°C using GentleMACS program TK3. The digested tissue was then filtered through a 100 μm cell strainer and washed with cold R-10. Samples were centrifuged at 500 *g* x 5 minutes at 4°C and supernatant was aspirated. Pelleted cells were resuspended in 1mL of ammonium-chloride-potassium (ACK) lysing buffer for 3 minutes at room temperature and then washed with 5 mL of R-10. Samples were centrifuged again at 500 *g* x 5 minutes at 4°C and supernatant was aspirated, the cell pellet was resuspended in 1mL of RPMI plus 2 uL of 20 mg/mL DNase and incubated at room temperature for 5 minutes. After incubation 4 mL of R10 was added and cells were counted. Cells were spun again at 500 *g* x 5 minutes at 4°C and resuspended in 1 million/mL aliquots in BAMBANKER Serum-Free Cell Freezing Medium (Wako Chemicals) for cryopreservation at -80°C.

### Single cell immune profiling

Samples were batched for multiplexed combined 3’ single-cell RNA, 5’ T-cell receptor (TCR), and cell-surface epitope assessment (CITE-seq) (10x Genomics.) Selected samples in batches of no more than 30 samples, were thawed from -80°C in a 37°C water bath with gentle agitation, transferred to a 15 mL conical tube and then slowly diluted with 9mL of RPMI. Samples were then centrifuged at 200 rpm for 10 min at 4°C and the supernatant was aspirated. Each sample was resuspended in 50 uL of 1:400 dilution in PBS of Ghost Dye 510 (Tonbo) and transferred to a 96-well V-bottom plate, then incubated at room temperature in the dark for 10 minutes. Samples were washed with 100 uL of PBS, centrifuged at 700 *g* for 2 minutes at 4°C, and supernatant was discarded. Each sample was then resuspended in 25 uL of Human TruStain FcX (Biolegand) diluted 1:20 in PBS and incubated for 10 minutes in the dark on ice. Then an antibody pool of TotalSeq-C Human Universal Cocktail V1.0 (Biolegend), TotalSeqC anti-human Hashtag Antibodies (Biolegend) was added, and samples were incubated for 30 minutes on ice. Samples were washed with 100 uL of PBS, centrifuged at 700 *g* for 2 minutes at 4°C and supernatant was discarded. Cells were then resuspended in 200 uL of 0.5% bovine serum albumin (BSA) in PBS and transferred to a flow tube with strainer cap. Cells were flow sorted for live cells and then pooled and counted. All samples were concentrated to a volume of 50 uL and then delivered to the Single Cell Analytics Innovation Lab (SAIL) for library construction and sequencing.

Single-cell RNA-Seq of FACS-sorted cell suspensions was performed on Chromium instrument (10x genomics) following the user manual for 5′ v2 chemistry. In brief, FACS-sorted cells were washed once with PBS containing 2% bovine serum albumin (BSA) and resuspended in PBS containing 2% BSA to a final concentration of 700–1,200 cells per μL. The viability of cells was usually above 70%, as confirmed with 0.2% (w/v) Trypan Blue staining (Countess II). Between 10,000 and 30,000 cells were targeted for each sample, where several samples were multiplexed together on one lane of 10x Chromium and specific extracellular protein epitope characterized (using Antibody Derived Tag - ADT) according to the cell hashing protocol.^7^

The scRNA-seq and scTCR-seq libraries were prepared using the 10x Single Cell Immune Profiling Solution Kit, according to the manufacturer’s instructions. Briefly, amplified cDNA was used for both 5′ gene expression library construction and TCR enrichment. For gene expression library construction, amplified cDNA was fragmented and end-repaired, double-sided size-selected with SPRIselect beads, PCR-amplified with sample indexing primers (98 °C for 45 s; 14–16 cycles of 98 °C for 20 s, 54 °C for 30 s, 72 °C for 20 s; 72 °C for 1 min), and double-sided size-selected with SPRIselect beads.

For TCR library construction, TCR transcripts were enriched from 2 μl of amplified cDNA by PCR (primer sets 1 and 2: 98 °C for 45 s; 10 cycles of 98 °C for 20 s, 67 °C for 30 s, 72 °C for 1 min; 72 °C for 1 min). Following TCR enrichment, enriched PCR product was fragmented and end-repaired, size-selected with SPRIselect beads, PCR-amplified with sample-indexing primers (98 °C for 45 s; 9 cycles of 98 °C for 20 s, 54 °C for 30 s, 72 °C for 20 s; 72 °C for 1 min), and size-selected with SPRIselect beads.

Final libraries (GEX, TCR and HTO/CITE) were sequenced on Illumina NovaSeq S4 or X+ platform (R1 – 26 cycles, i7 – 10 cycles, i5 – 10 cycles, R2 – 90 cycles).

### Demultiplexing

HTO sequencing data were aligned to the HTO barcodes, and UMIs were counted for each cell using CITE-seq-Count. Using a two-component K-means algorithm, we partitioned logged HTO counts into two distributions: background noise (lower mean) and positive tags (larger mean). Each droplet was then assigned to its source sample based on tags in the positive signal component. We classified droplets with multiple assignments as doublets and those with a single assignment as singlets. This analysis was performed using SHARP v0.1.1 (https://github.com/hisplan/sharp)^8^.

### Single-Cell Analysis

We used a hierarchical supervised approach to assign phenotypes to cells as follows. After QC and pre-processing, we used unsupervised graph-based clustering to define the main CD45+ cell types. This yielded 6 separate clusters, namely T/NK, B, Plasma, Mast, Myeloid, and Plasmacytoid DC (Figure 2a), annotated using known marker genes (Extended Fig. 1, Supplementary Table 4).

### Preprocessing

Raw binary base call (BCL) files were demultiplexed into FASTQ files using Cell Ranger. Gene expression libraries were aligned to the GRCh38 reference genome and quantified using `cellranger count` (v7.1.0). TCR V(D)J libraries were processed with `cellranger vdj` (v6.1.2). Cell barcodes were selected using Cell Ranger’s cell-calling algorithm EmptyDrops, and the resulting whitelist was used to filter ADT data. The ADT data was then quantified using the SHARP pipeline, which builds on CITE-seq-Count to count cell barcodes and UMIs. Finally, these matrices—gene expression, ADT, and TCR—were loaded into individual AnnData objects for downstream single-cell analysis.

Cells with fewer than 200 genes and more than 40% mitochondrial content were removed and samples with fewer than 100 cells were removed. Doublet cells were identified using scDblFinder^9^ and excluded. Raw counts were normalized by library size and log-transformed with a pseudo-count of 1 using Scanpy.^10^ The top 2000 highly variable genes were selected following the removal of ribosomal, mitochondrial and T cell receptor genes using the `highly_variable_genes` function with the `flavor` option set to `seurat` and sample as `batch_key`. Principal component analysis (PCA) was performed retaining the top 50 PCs. `Harmonypy`^11^ (v0.0.9) was used to perform batch correction to control for inter-patient variation. The batch corrected embeddings were used for leiden clustering and creating the UMAP^12^ projection for visualization. Clusters for the broad cell lineages were annotated using marker genes, derived from Wilcoxon rank-sum tests comparing each cluster to all others. The 8 annotated clusters were: B (*MS4A1, CD19, CD79A*), T/NK (*CD3D, CD3E, NCAM1*), epithelial (*EPCAM, KRT19, KRT8*), endothelial (*PECAM1, VWF, CDH5*), fibroblasts (*COL1A1, FAP, PDGFRA*), myeloid (*CD14, LYZ, CSF1R*), mast (*KIT, TPSB2, FCER1A, CPA3, HDC, MS4A2*), and plasma (*MZB1, SDC1, JCHAIN*). Using these annotated cell type clusters, the data were subset to CD45+ cells and cells with more than 10% mitochondrial content were removed. Highly variable gene selection, PCA, batch correction, leiden clustering and UMAP computation were performed as outlined above.

### Cell-type annotations

CD45+ cells were annotated using a combination of gene expression and protein marker intensity. Cells were grouped into major compartments, namely T/NK, myeloid, B, plasma, mast and plasmacytoid DC cells (**Supplementary Table 4**). T cells were annotated using cite-seq protein CD3D, with custom intensity thresholds to filter for CD4+ and CD8+ T cells using CD8A and CD4 proteins, respectively. For CD4, CD8 and myeloid cells, clustering analysis was performed and the UMAP re-computed for each subset. Differentially expressed genes between clusters were evaluated alongside protein expression (e.g. CD45RA and CD45RO) to annotate 13 CD8, 11 CD4 and 7 myeloid subclusters (**Supplementary Table 4**).

### Gaussian Kernel Density Estimates

We estimated cell density across the UMAP embedding using Scanpy’s `sc.tl.embedding_density` with default parameters, grouping cells by the respective covariate. For each group, we visualized the resulting Gaussian kernel density using `sc.pl.embedding_density.`

### Tensor factorization for Cytokine Activation using IREA

Our objective was to identify intercellular cytokine driven signaling bridging tumors and tumor-draining lymph nodes. To do so, we computed a transcriptional impact for each of a list of 86 cytokines in each cell type per sample. To account for the variation of cell types and cytokines in our samples, we used a Bayesian Poisson tensor factorization algorithm shown to be robust to sparse datasets.^13^ The algorithm infers sample-level representations in terms of activities of latent multi-cytokine factors, which represent a group of cytokines that coordinately impact multiple cell types across samples. For each anatomical site of origin, we grouped samples based on their factor activation scores, using a nearest-neighbor graph community detection algorithm.

Concretely, top up-regulated genes in each cell type for each cytokine were acquired from a recently published cytokine dictionary (hereafter called IREA genes.)^14^ For each sample, and each cytokine, the overlap between the IREA genes and the DE genes in each cell type was used to compute an overrepresentation analysis and p-values were computed via the hypergeometric test and corrected for multiple testing using false discovery rate (FDR). An enrichment score was computed as the negative log10 of the adjusted p-values per (sample, cell-type, cytokine) tuple. A sample by cytokine score was computed as the sum of all cytokine-cell type interactions. The sample by cell type by cytokine enrichment score, computed thus, was used as the input to the tensor factorization algorithm. Let 𝑥_𝑖𝑗𝑐_ denote the enrichment of cytokine 𝑐 , in cell type 𝑗 , in sample 𝑖 , then we assume the following decomposition: 𝑥_𝑖𝑗𝑐_ ≈ 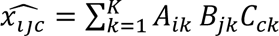 where 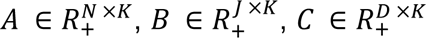 are latent non-negative matrices, representing sample by factor, cell type by factor, and cytokine by factor scores respectively. To encourage sparsity in the latent factors, we placed Gamma priors on each factor 𝐴_𝑖𝑘_ ∼ Gamma (𝛼, 𝛽) , 𝐵_𝑗𝑘_ ∼ Gamma (𝛼, 𝛽) , and 𝐶_𝑐𝑘_ ∼ Gamma (𝛼, 𝛽) , and adopted a Poisson likelihood 𝑥_𝑖𝑗𝑐_ ∼ Poisson( . 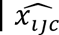). The Poisson likelihood favors zero values for small rates, and its closed-form probability mass function accommodates non-negative real-valued inputs.^15^ The target of inference is the posterior distribution on the latent matrices 𝐴 , 𝐵 , 𝐶 , that is 𝑃( 𝐴, 𝐵, 𝐶 ∣ 𝑋, 𝜆 ) . We used variational inference to infer the latent variables.^16^

### Cell type-Cytokine Activation Graph Construction

Let 𝜓_𝑖𝑗_ denote the activity of cytokine 𝑖 in cell type 𝑗 . We used 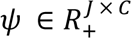 as a weighted adjacency matrix for a bipartite graph where nodes are the set of cytokines and cell types, and edges denote the inferred activity level of a source cytokine in a receiver cell type. To compute 𝜓, we averaged over factors to compute sample-specific (cell-type, cytokine) pairings, for each sample, per cell type per cytokine 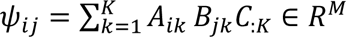. For each group of samples 𝐺 , we computed a cell type-cytokine activation matrix as 𝜓_𝐺𝑗_ ≔ ∑_𝑖∈𝐺_ 𝜓_𝑖𝑗_ . Similarly, for a set of cytokines 𝐻 , we reported a summary 𝜓_𝐺𝐻_ as 𝜓_𝐺𝐻_ ≔ ∑_𝑖∈𝐺,𝑗∈𝐻_ 𝜓_𝑖𝑗_.

### Cytokine type groupings

We grouped into biologically relevant cytokine families as below:

- Class 2 cytokines: IL-10 and all the interferons (k, g, e, b, a1)
- Common beta-chain cytokines: IL3, IL5, GM-CSF
- Colony stimulating factors: G-CSF, M-CSF, Flt3l
- Hormones: Prolactin, Adiponectin
- IL-1/Alarmin family: TSLP, IL1a, Il1b, IL18, IL33, IL36a
- IL-12 family: IL12, IL23, IL27
- IL-17 family: IL17E
- IL-2 family: IL2, IL7, IL15, IL21
- IL-6 family: IL11, IL31, LIF, Cardiotrophin, Neuropoietin, Oncostatin M
- Th2-like: IL4, IL9, IL13
- TNF family: TNF, TL1A, CD40L, BAFF

### Clustering of samples based on IREA scores

We grouped samples that belong to each anatomical site separately. Briefly, given the sample by factor matrix 𝐴 , we selected rows corresponding to samples from the dLN (tumor), formed the nearest neighbor graph and computed the leiden clustering with resolution 0.5. For tumor samples, this initially yielded 4 clusters. For interpretability, we merged the two that were high in T cell interactions.

### Receptor Ligand activation analysis

We inferred cell-cell interactions using CellChat,^17^ calculating communication probabilities for each cell type pair and associated significance using permutation testing for disrupted and canonical lymph node samples separately. We merged these CellChat objects to perform comparisons between the two groups and to identify significantly differing ligand-receptor interaction probabilities. We then filtered these results to cytokine interactions. To further evaluate the evidence for these predicted interactions, we compared the gene expression for each significant ligand or receptor highlighted by CellChat. This was performed by pooling all cells from each lymph node group and comparing gene expression using a Wilcoxon rank-sum test with Benjamini–Hochberg correction for multiple comparisons. This strategy allowed us to include all samples (particularly valuable for myeloid cell types due to the relatively low cell counts for this cell lineage). Gene expression after pseudobulking by sample was assessed and only genes that showed a consistent trend between the pseudobulked and pooled data were retained in our results.

### CD8+ T cell Trajectory Analysis

For trajectory analysis, we used Palantir and CellRank. First, we selected CD8+ T cells derived from dLN and tumor samples based on prior annotations. We then identified the top 2,000 highly variable genes using the Scanpy function `sc.pp.highly_variable_genes`. Next, we computed the top 50 principal components using `sc.tl.pca` and performed batch correction via Harmony integration across individual patient identifiers using `sc.external.pp.harmony_integrate`. We computed a diffusion map based on these Harmony-corrected PCA embeddings using `sc.tl.diffmap` to capture global structure and lineage relationships. We initialized the pseudotime trajectory by selecting a root cell representative of a biologically meaningful starting point—specifically, a cluster of phenotypically naïve T cells identified using our annotations. We then used Palantir’s `palantir.core.run_palantir` function to infer cell fate potential and generate pseudotime values, employing 1,000 waypoints. Finally, we use CellRank’s `cellrank.tl.terminal_states` and `cellrank.tl.lineages` functions, leveraging Palantir-derived pseudotime and diffusion-based transition probabilities to identify terminal states and major cellular lineages. We executed CellRank with 1,000 permutations to identify lineage drivers and associated genes.

### Phenotype entropy

We first compute the fraction of CD8+ T cells per sample. We then compute the Shannon entropy

as 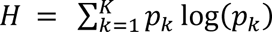.

### Identifying naïve cell subpopulations

We used Gaussian Mixture Models (GMM) to identify subpopulations of naïve cells based on the protein expression of CD45 isoforms. Briefly, we selected all cells that had a naïve phenotype based on their transcription profile. We used Bayesian information criterion to select the number of components for the GMM. We used the from `sklearn.mixture.GaussianMixture` with a full covariance, `max_iter=10000`, and 100 random restarts.

### Distance between T Cell Phenotypes

To compute the distance between different T cell phenotype clusters, we used the geometric median computed Weiszfeld’s algorithm.^18^ Geometric median of a set of 𝑛 points {𝑋_𝑖_}, is defined as the point 𝑦 that minimizes the sum of distances from all poits in the set, i.e., 𝑦 ≔ argmin*_𝑦∈𝑅_^D^* 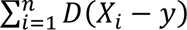, where 𝐷 is the 𝐿_2_ norm (Euclidean distance). We used the diffusion map representation of cells as input. We computed the geometric median for each phenotype group the computed the pairwise distance of geometric medians of each phenotype cluster from all others.

### Tumor vs dLN Transcription Factor Comparison

To find which transcription factors (TF) affect expansion in CD8+ T cells, we compared the expression of TFs in CD8+ T cells, excluding the naïve phenotype, derived from Tumor to draining lymph node. We obtained a list of human TFs^19^ and ranked their expression using the Wilcoxon rank-sum test.

### KLF2 Correlated Genes

We used regularized linear regression to find genes predictive of KLF2’s expression in a sample specific manner. Gene expression counts were normalized for library-size, log-transformed, and standardized such that the mean and the variance of each gene are zero and one respectively. Using linear regression with an L2 penalty (i.e., ridge regression), we estimated a per sample coefficient vector of per gene association with KLF2 expression. We used the collection of these coefficients to compute a mean coefficient and Z-score per gene.

### KLF2 Gene Cloning

The plasmid PpyCAG-KLF2-IB^20^ was a gift from Austin Smith (Addgene #60441), which contained human *KLF2*. PCR amplification of *KLF2* was performed, using primers (in table below) and Q5® High-Fidelity DNA Polymerase (New England Biolabs) according to manufacturer’s instructions. A restriction digest was performed of both source plasmid PpyCAG-KLF2-IB and destination plasmid pTRIP-SFFV-mTagBFP-P2A (Addgene #102585), using BamHI-HF (New England Biolabs) and SalI-HF (New England Biolabs) with rCutSmart™ Buffer (New England Biolabs) according to manufacturer’s instructions, gel isolated and purified. Construct assembly was then performed using Gibson Assembly® Master Mix (New England Biolabs) according to manufacturer’s instructions. The constructed plasmid was then transformed into NEB 5-alpha competent *E. coli* cells (New England Biolabs) using 1uL of plasmid DNA. Mini-prep using Qiagen Mini Kit (Qiagen) was performed for 3 individual clones per manufacturer’s instructions, and then Sanger sequencing was used to confirm the appropriate sequence for clone 2 which was subsequently used for all experiments.

### Cloning primer sequences

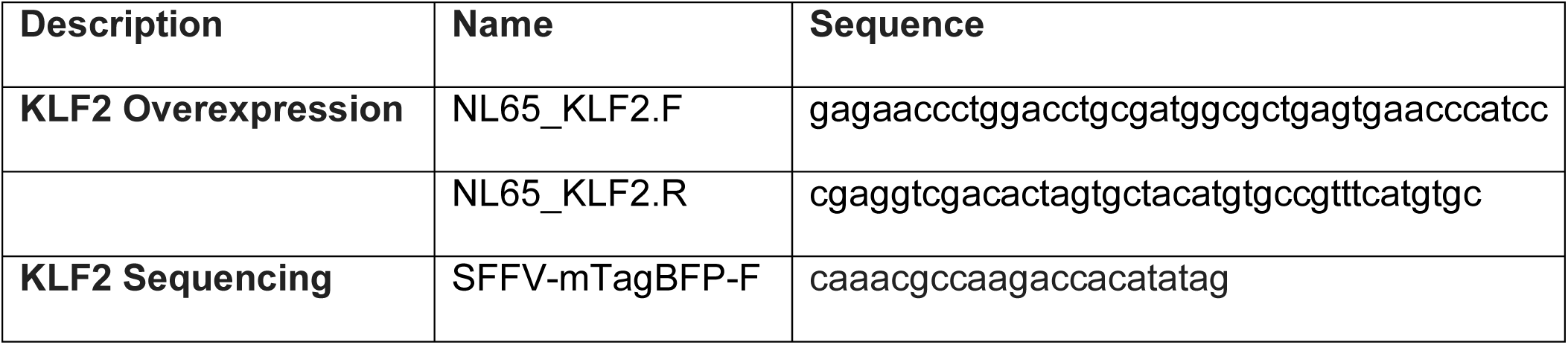

### KLF2 Overexpression

Lentivirus was produced using 293T cells as follows: briefly, 293T cells were transiently co-transfected with viral packaging (pCG-gag-pol), and envelope (pCG-VSVg) constructs along with the expression vector (pTRIP-SFFV-mTagBFP-P2A-Empty or pTRIP-SFFV-mTagBFP-P2A-KLF2) using 100 mg/mL polyethylenimine (Polysciences Inc). Virus-containing supernatant was collected beginning 48 hours after transfection and filtered with a 0.45µm syringe, then aliquoted and stored at -80°C until use. Primary human T-cell activation was performed as follows: Buffy coats containing PBMCs were obtained from New York Blood Center. PBMCs were isolated using Lymphoprep density gradient-based centrifugation. Subsequently, T cells were isolated using a Dynabeads Untouched Human T Cell kit according to manufacturer’s instructions. T cells were activated in T cell media (RPMI-1640 supplemented with 10% heat-inactivated FBS, 2 mM *L*-glutamine, and 100 U/mL Penicillin-Streptomycin) supplemented with recombinant IL-2 (Peprotech, 5 ng/mL). Cells were plated onto non-tissue culture-treated 6-well plates that had been coated with anti-CD3 (Biolegend, 5 µg/mL) and anti-CD28 antibody (Biolegend, 2 µg/mL) in PBS. 24 hours after activation, 100x concentrated virus was diluted 1:50 in T cell media (40 uL virus in 160 uL T cell media, see above), then 200 uL was added dropwise to each well and plates were shaken gently. 24 hours later, cells were passaged into tissue culture-treated 6-well plates containing fresh T cell media supplemented with IL-2. 5 days following initial stimulation, cells were either harvested directly for spectral cytometry-based phenotyping, DOGMA-Seq, assessment of KLF2 by Western blot, or restimulated to interrogate intracellular cytokine production as described below.

### Protein extraction and Western Blot

Cell lysis was performed using RIPA buffer (Cell Signaling) containing 1% SDS as well as protease and phosphatase inhibitors. Following lysis, cells were sonicated (Bioruptor) and centrifuged and supernatants were analyzed for protein concentration using BCA. 40 ug of each lysate were run on a 4-12% Bis-Tris Gel (Thermo Fisher) and transferred to nitrocellulose membranes. Membranes were blocked using 5% milk and then blotted with the following antibodies: anti-KLF2 (see table below) and anti-H3 (see table below) in 3% BSA overnight at 4C on a rocker. Membranes were washed and then incubated with HRP-conjugated secondary antibodies at 1:5000 dilution for 1 hour at room temperature in 5% milk. Membranes were then washed and imaged using a Bio-Rad ChemiDoc Imaging System.

| Target Name | Host | Company | Catalog # | Dilution |
| --- | --- | --- | --- | --- |
| KLF2 | Mouse | Novus Biologicals | NBP2-61812 | 1:1000 |
| Histone H3 | Rabbit | Cell Signaling | 9715 | 1:1000 |
| Anti-mouse IgG, HRP-linked Antibody | Horse | Cell Signaling | 7076 | 1:5000 |
| Amersham ECL Rabbit IgG, HRP-linked whole Ab | Donkey | Cytiva | NA934 | 1:5000 |

### Spectral Cytometry

For phenotyping, cells were incubated with Fc block (Human TruStain FcX, Biolegend, cat. 422302) at a 1:20 dilution for 30 minutes at 4°C. Viability staining was performed for 30 minutes at 4°C using Ghost Dye 510 Violet (cat. 59863) diluted 1:800 in PBS. Cells were subsequently washed and stained for surface markers in FACS buffer containing 1X Brilliant Buffer (cat. 566349) for 30 minutes at 4°C, then washed twice and resuspended in 1% BSA in PBS. Data was acquired using a Cytek Aurora spectral cytometer.

| Staining type | Marker | Fluorophore | Catalog # | Clone | Dilution |
| --- | --- | --- | --- | --- | --- |
| Phenotype, Surface | PD-1 | APC | BioLegend, 329908 | EH12.2H7 | 1:100 |
|  | CCR7 | BB700 | BD Biosciences, 566437 | 3D12 | 1:50 |
|  | CD45RA | BUV395 | BD Biosciences, 740298 | HI100 | 1:400 |
|  | CD8 | BUV496 | BD Biosciences, 612943 | RPA-T8 | 1:800 |
|  | CD38 | BUV661 | BD Biosciences, 612969 | HIT2 | 1:200 |
|  | HLA-DR | BV786 | BD Biosciences, 564041 | G46-6 | 1:200 |
|  | CD4 | BV570 | BioLegend, 300534 | RPA-T4 | 1:800 |
|  | CD28 | BUV805 | BD Biosciences, 742037 | CD28.2 (RUO) | 1:100 |
|  | CD69 | BV605 | BioLegend, 310937 | FN50 | 1:200 |
|  | Tim-3 | BV650 | BioLegend, 345028 | F38-2E2 | 1:50 |
|  | CD103 | PE-594 | BioLegend, 350223 | Ber-ACT8 | 1:100 |
|  | CD39 | PE-810 | BioLegend, 328245 | A1 | 1:100 |
|  | CXCR3 | A700 | BD Biosciences, 561320 | 1C6/CXC R3 | 1:100 |
|  | CXCR6 | BV421 | BioLegend, 356013 | K041E5 | 1:100 |
| **Phenotype, Intracellular** | TCF-1 | A488 | Cell Signaling, 6444S | C63D9 | 1:100 |
|  | TOX | PE | MACS Miltenyi Biotec, 130-120-716 | REA473 | 1: 25 |
|  | CTLA4 | BV711 | BioLegend, 369632 | BNI3 | 1: 200 |

For interrogation of cytokine production, PMA and ionomycin were added to wells to a final concentration of 50 ng/mL and 500 ng/mL, respectively. 1 hour after addition of PMA and ionomycin, Brefeldin A was added at a 1:1000 dilution and cells were incubated for another 3 hours, after which they were harvested and stained with surface antibodies as described above using the panel below. Fixation was performed using an eBioscience FoxP3 Fixation/Permeabilization kit and stained overnight with intracellular antibodies diluted in permeabilization buffer overnight at 4C. Data was acquired using a Cytek Aurora spectral cytometer.

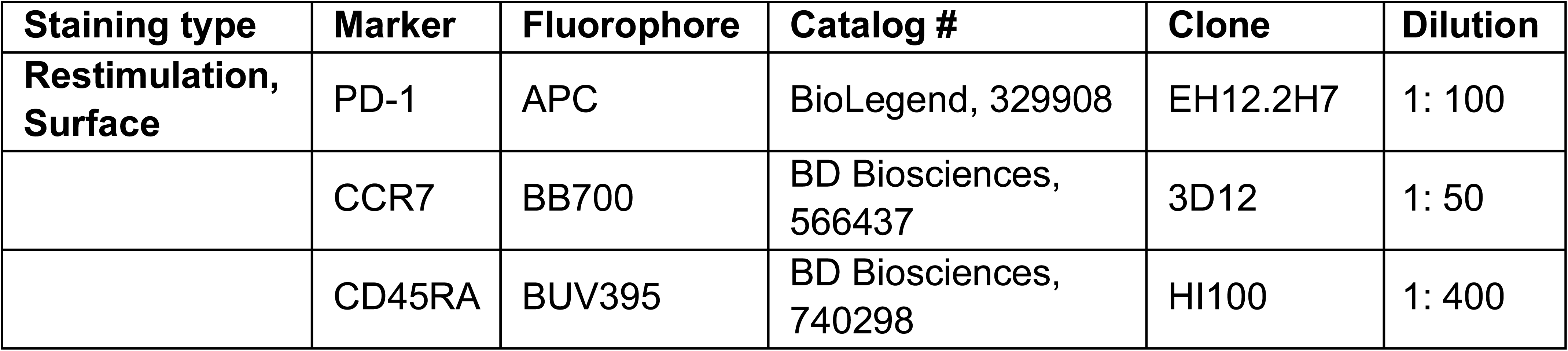
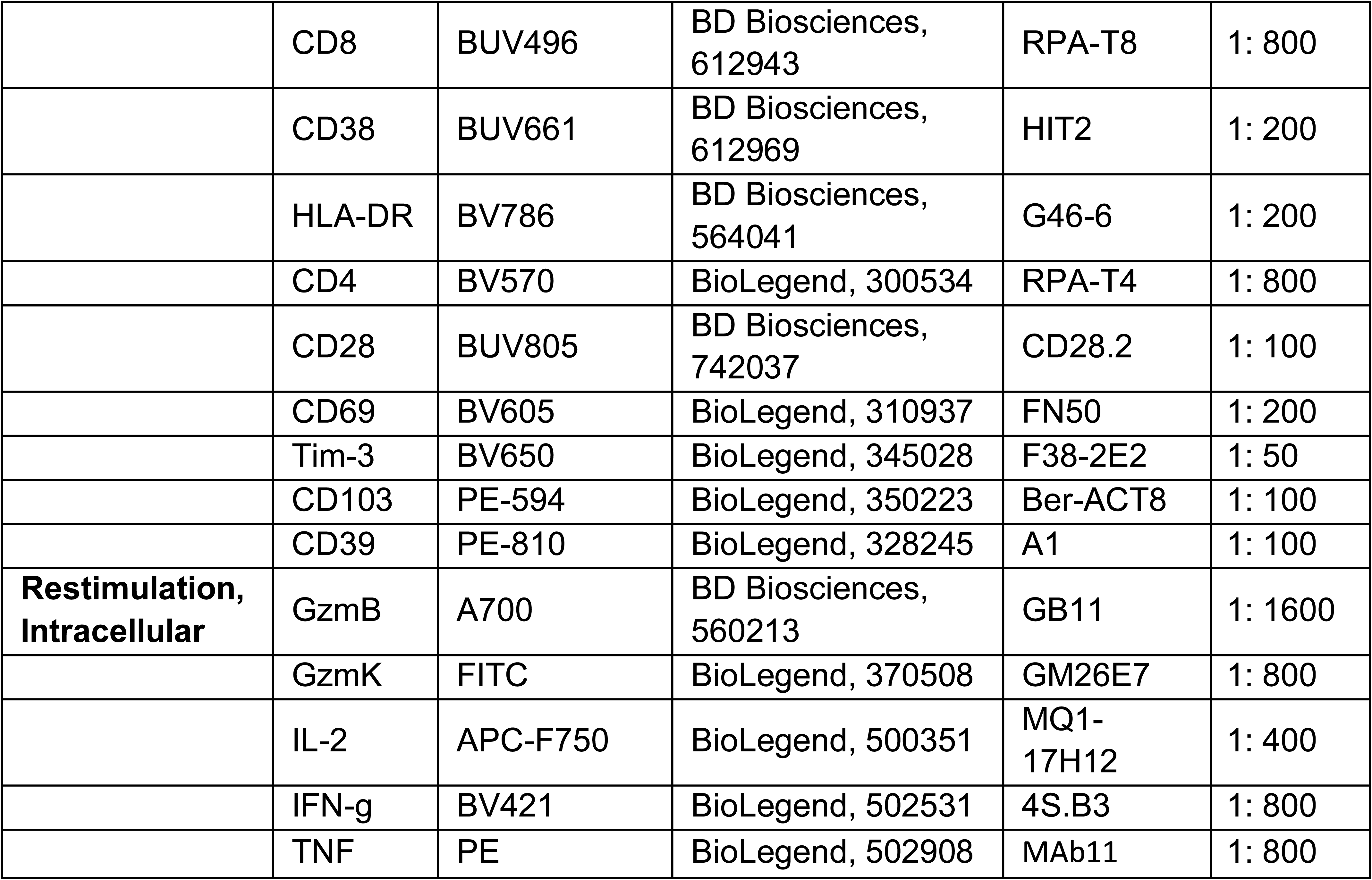

### DOGMA-Seq

Primary human T cells were isolated and cultured as described above (‘**KLF2 Overexpression**’). On day 5 following initial activation, T cells were collected and stained with a viability dye followed by surface staining with anti-CD4 (RPA-T4, BioLegend 300534, 1:100), anti-CD8 (RPA-T8, BD Biosciences 612914, 1:1200), TotalSeq-A Human Universal Cocktail, and TSA-compatible hashtag oligos for multiplexing. Live BFP+ CD4^+^ and CD8^+^ T cells from both KLF2 overexpressing and empty vector conditions were flow sorted and delivered to SAIL for sequencing. Single Cell Multiome ATAC + Gene Expression was performed with 10x genomics system using Chromium Next GEM Single Cell Multiome Reagent Kit. We followed User Guide with the difference that single cell suspension was prepared as described in Mimitou et al (DOGMA-seq protocol with Digitonin buffer).^21^ Around 10,000 cells were targeted for each sample. Samples were multiplexed together on one lane of 10x Chromium (using Hash Tag Oligonucleotides - HTO) and specific extracellular protein epitope characterized (using Antibody Derived Tag - ADT) following previously published protocol.^7^ Next, cells were treated with a digitonin lysis buffer (20 mM Tris-HCl pH 7.4, 150 mM NaCl, 3 mM MgCl2, 0.01% DIG and 2 U µl−1 of RNase inhibitor) for 5 min on ice, followed by adding 1 ml of chilled DIG wash buffer (20 mM Tris-HCl pH 7.4, 150 mM NaCl, 3 mM MgCl2 and 1 U µl−1 of RNAse inhibitor) and inversion before centrifugation at 500g for 5 min at 4 °C. The supernatant was discarded, and cells were resuspended in DIG wash buffer, followed by counting using trypan blue and DAPI on a Countess II FL Automated Cell Counter. Then cells were processed according to the Chromium Next GEM Single Cell Multiome ATAC + Gene Expression user guide. Final libraries were sequenced on Illumina NovaSeq X+ platform following 10x genomics recommendations.

For sample demultiplexing, HTO sequencing data were aligned to the HTO barcodes, and UMIs were counted for each cell using CITE-seq-Count. Using a two-component K-means algorithm, we partitioned logged HTO counts into two distributions: background noise (lower mean) and positive tags (larger mean). Each droplet was then assigned to its source sample based on tags in the positive signal component. We classified droplets with multiple assignments as doublets and those with a single assignment as singlets. This analysis was performed using SHARP v0.1.1 (https://github.com/hisplan/sharp)^8^.

### Cell typing of DOGMA-seq

The KLF2 over-expression and empty vector were demultiplexed as above in ‘**DOGMA-Seq**’. We processed single-cell RNA-, ATAC- and CITE-seq (DOGMA-seq) data using the Seurat (v. 5.0.3) and Signac packages (v.1.13.0). A total of 9,616 cells were sequenced, 6,487 in the KLF2 over-expression and 3,129 in the empty vector conditions. We loaded RNA and ATAC-seq data into a single Seurat object. We computed quality control metrics—including nucleosome signal and transcription start site (TSS) enrichment and filtered cells based on RNA/ATAC count thresholds, nucleosome signal, and TSS enrichment to exclude low-quality cells. Next, we established a unified peak set by merging and filtering called peaks, removing non-standard chromosomes and blacklist regions. We quantified ATAC counts within these peaks using `FeatureMatrix` and added them to a new chromatin assay in the Seurat object. We then split the CD4+ and CD8+ T cells based on the CD4 and CD8A protein expression. To annotate cell phenotypes, we normalized the RNA expression counts with `SCTransform` and reduced dimensions using `RunPCA`. We processed ATAC data with term frequency-inverse document frequency (`RunTFIDF`) and `RunSVD`. We then integrated RNA and ATAC modalities using `FindMultiModalNeighbors` and `FindClusters` on the resulting joint weighted nearest-neighbor graph to identify clusters. We used `FindAllMarkers` on the SCT assay and separately on the ADT assay to report marker genes and proteins per cluster. We used these markers to annotate the T cell phenotypes.

### Motif Accessibility Analysis

We used the `RunChromVAR` function with default parameters from the Signac package to compute transcription factor (motif activities from single-cell chromatin accessibility data. Briefly, it quantifies how accessible each cell is at genomic regions associated with specific TF motifs by comparing observed accessibility to background expectations. ChromVAR generates bias-corrected, normalized activity scores (z-scores) for each motif in each cell, enabling the identification of TFs that are potentially driving differences in chromatin states across cell populations or experimental conditions. With the accessibility scores at hand, we used the `FindMarkers` function to compute the differentially accessible markers per cell type (with parameters `mean.fxn = rowMeans` and `fc.name = “avg_diff”`).

### KLF2 Targets

We first extracted KLF2’s motif’s binding sites that overlapped with any identified peak. We used the motif MA1515.2.^22^ We then filtered for regions that were within 1000bp of a protein coding gene’s promoter to identify putative target genes. Among these genes, we reported those with a log-fold-change (between the KLF2 over expression versus the empty vector conditions) of above 1.0 or below –1.0 and a gene expression value of 0.1 in at least one of the conditions.

### LFA-1 dependent T cell priming assay

#### Cell Isolation

OT-I T cells and dendritic cells were isolated as follows: for DC isolation, spleens from C57/Bl6 mice were minced and incubated in digestion buffer (200X DNase, 50X Liberase, RPMI) for 30 minutes at 37 degrees. During this incubation period, spleens from OT-I mice were isolated, mashed over a 40 µm strainer. Subsequently, spleens were washed with RPMI and treated with hypotonic ACK lysis buffer to lyse red blood cells. RBC lysis was quenched with 10x excess PBS and centrifuged. OT-I T cells and DCs were then enriched using T-cell isolation (Dynabeads™ Untouched™ Mouse T Cells Kit) and DC enrichment kits (Dynabeads™ Mouse DC Enrichment kit), respectively. These are both negative selection kits in which cell suspensions were first incubated with antibodies for 20 minutes at 4 degrees, subsequently incubated with magnetic beads for 15 minutes with gentle tilting and rotation, and finally placed on a magnet after which suspensions containing OT-I T cell and dendritic cell-enriched populations were transferred to a fresh tube for downstream assays.

#### T cell-DC coculture

After isolation, DCs were resuspended in complete RPMI-1640 media containing 10% FBS, 2 mM L-glutamine and Pen/Strep and incubated at 37 degrees for 1 hour with gentle rotation in the presence of escalating doses of SIINFEKL (N4), SIIQFEKL (Q4), SIITFEKL (T4), or SIIVFEKL (V4) ranging from 0.1, 1, 10, and 100 nM. Cells were subsequently washed and resuspended at 1 million cells/mL.

During this period, OT-I T cells were labeled with CellTrace Violet by resuspending in PBS at 1 million cells/mL, warming to 37 degrees, mixing 1:1 with CellTrace Violet diluted 1:1000 in pre-warmed PBS and incubating in a 37-degree water bath for 20 minutes. Labeling was quenched by adding 5x volume of RPMI+++. Labeled T cells were counted, centrifuged and resuspended in complete T cell media (RPMI+++ containing 50 uM BME and 10 ng/mL IL-2) at 200K/mL, with either isotype control (rat IgG2a kappa, BioXCell) or anti-LFA-1 (M17/4, BioXCell) at a final concentration of 5 ug/mL. 500 uL containing 100K T cells were added to each well of a 96 well Polypropylene Deep Well Plate (Corning) into which 5 uL DCs (5,000 DC total) were added. The plate was then incubated at 37 degrees for 4 days.

For interrogation of cytokine production, PMA and ionomycin were added to wells to a final concentration of 50 ng/mL and 500 ng/mL, respectively. 1 hour after addition of PMA and ionomycin, Brefeldin A was added at a 1:1000 dilution and cells were incubated for another 3 hours, after which they were harvested and stained with surface antibodies as described above using the panel below. Fixation was performed using an eBioscience FoxP3 Fixation/Permeabilization kit and stained overnight with intracellular antibodies diluted in permeabilization buffer overnight at 4C. Data was acquired using a Cytek Aurora spectral cytometer.

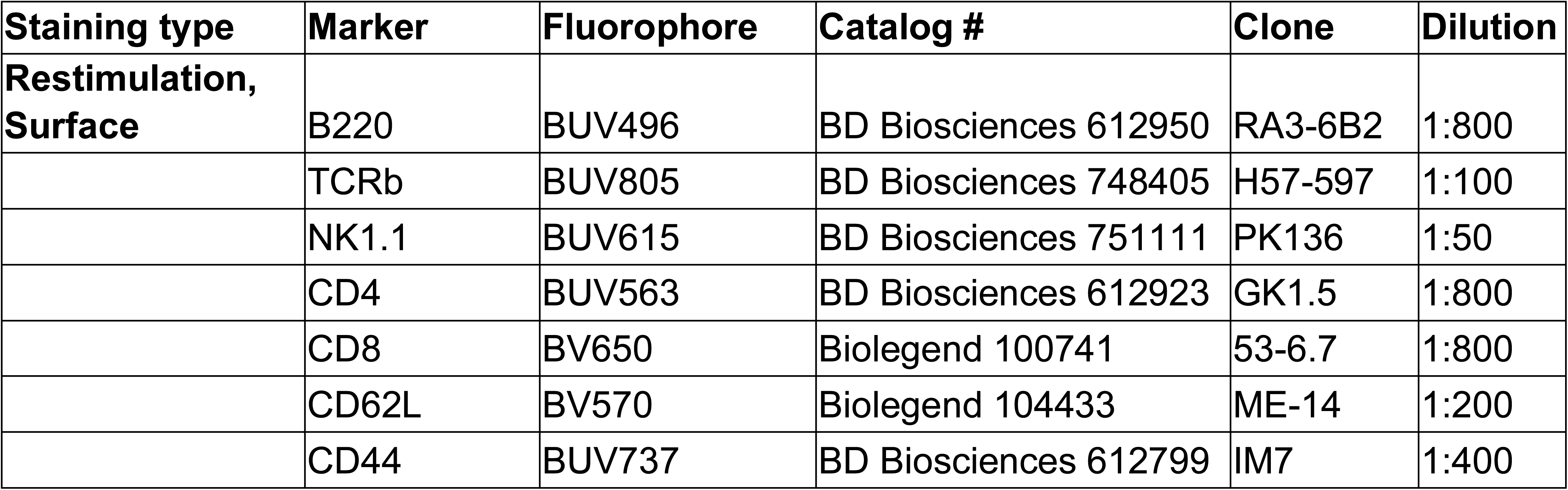
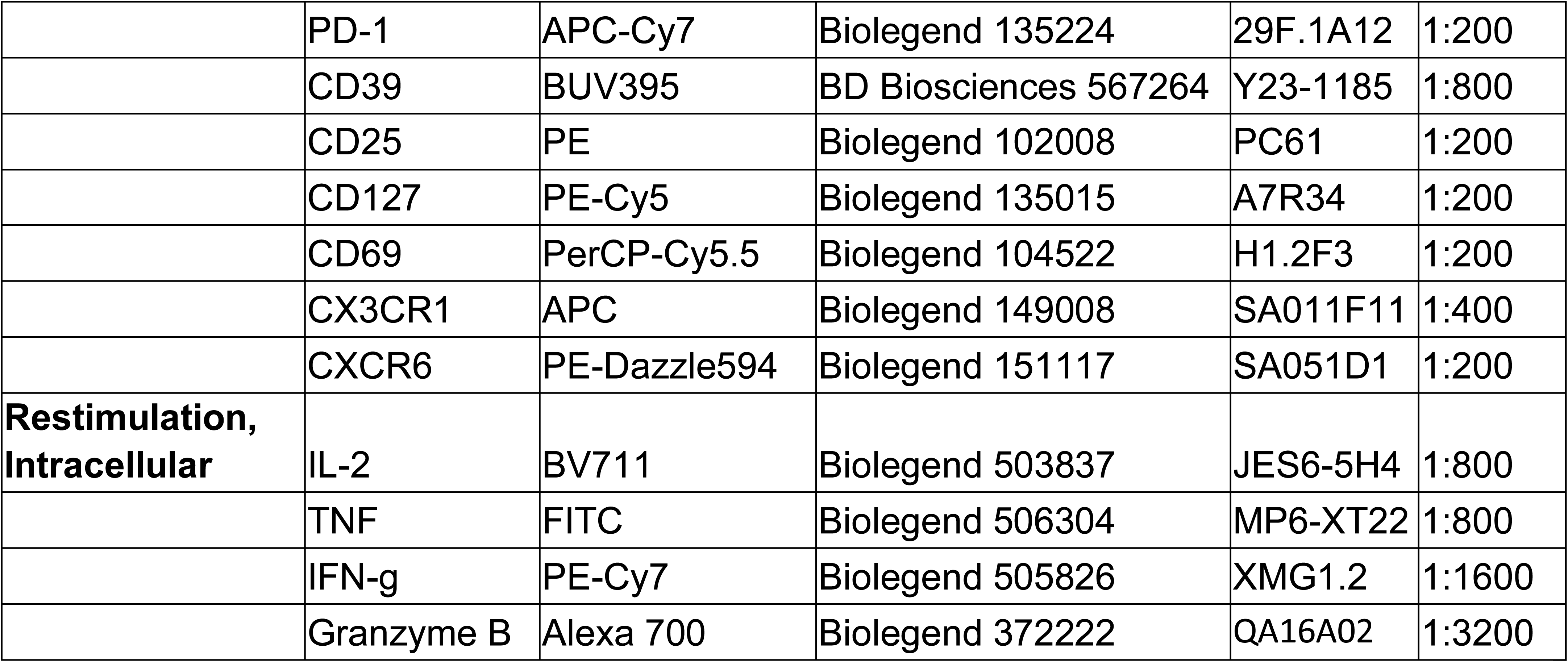

