## Extended Figures 1-7 for "Disrupted priming within draining lymph nodes drives immune quiescence in gastric cancer"

### Extended Figure 1

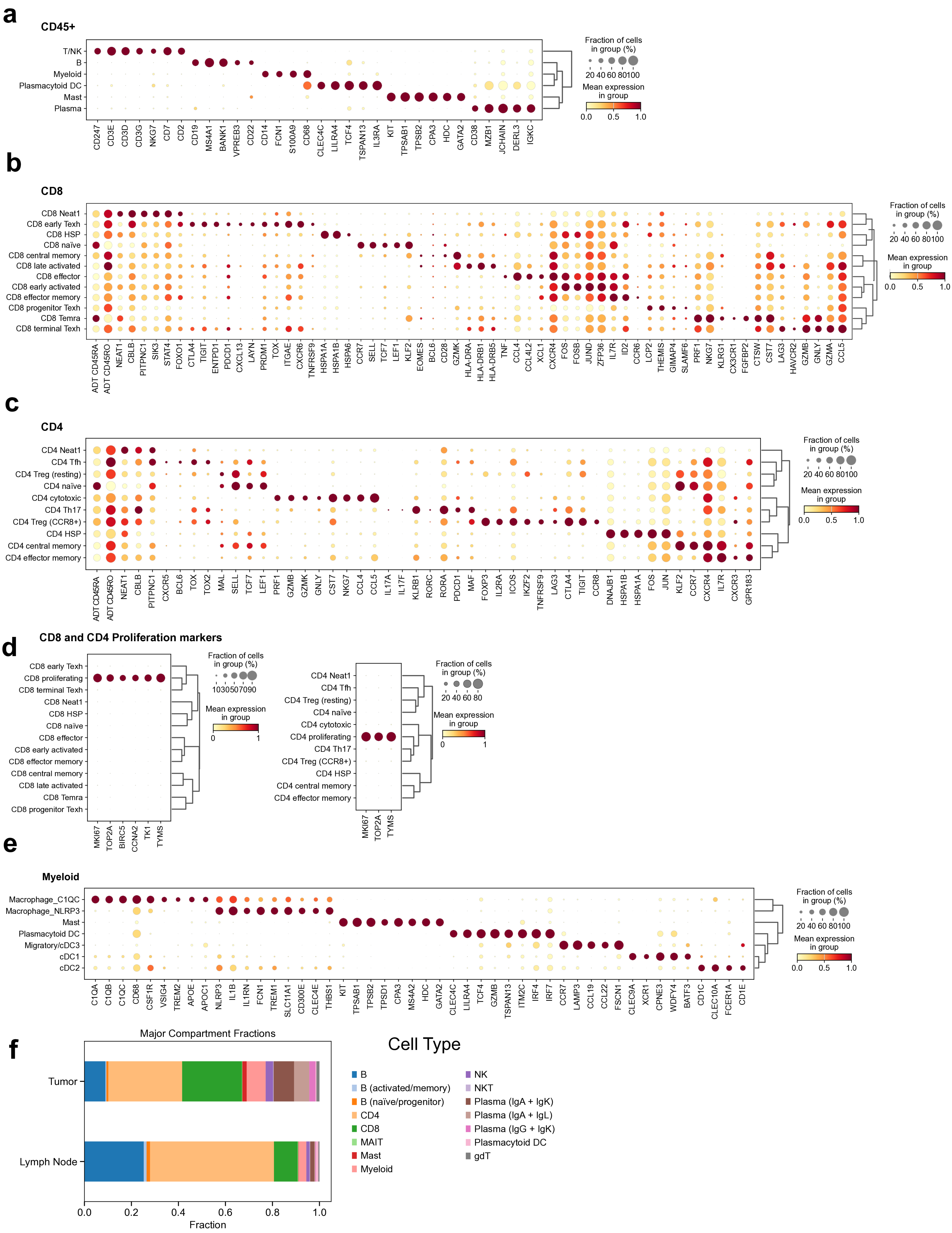

### Extended Figure 2

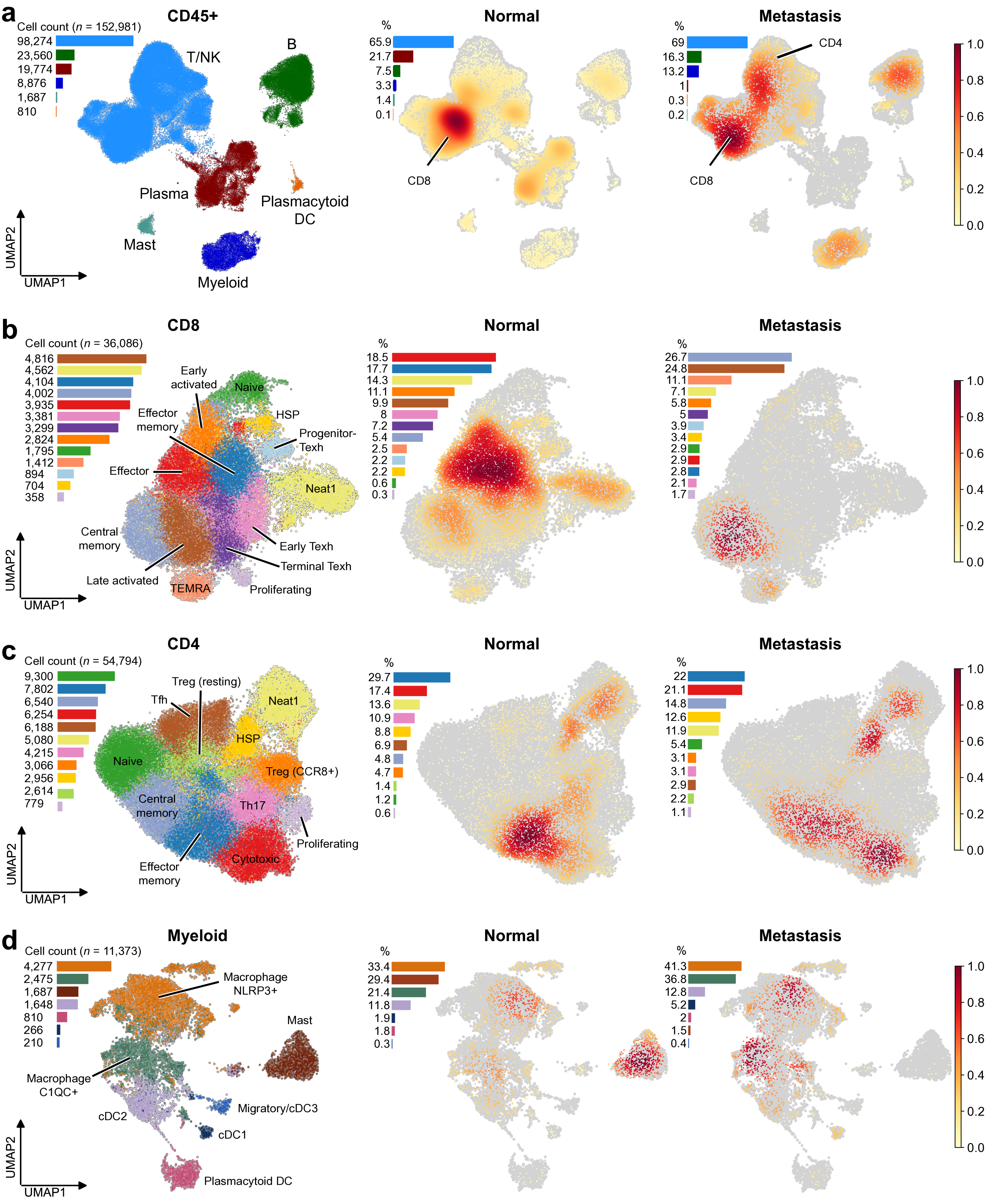

### Extended Figure 3

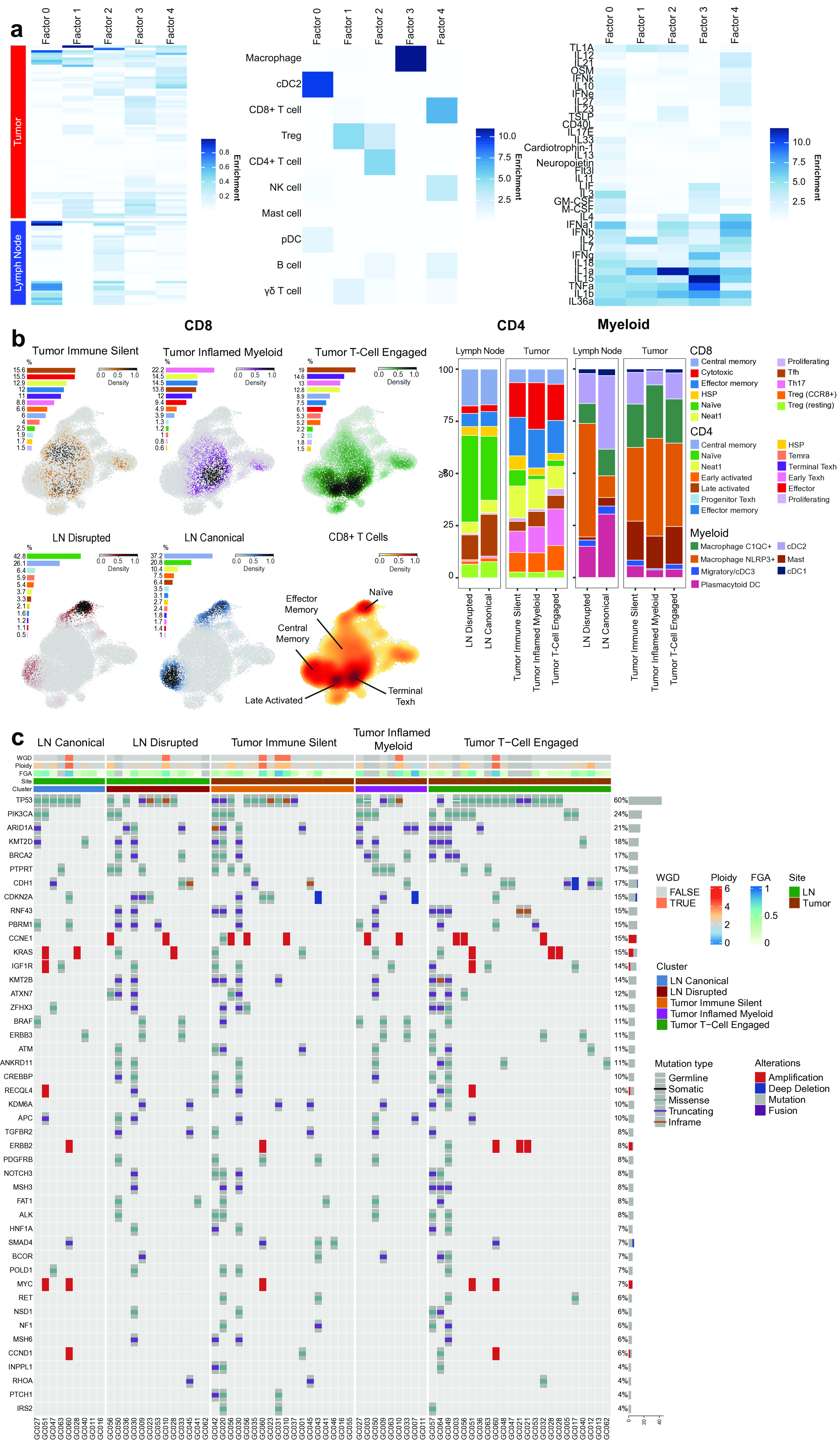

Extended Figure 4

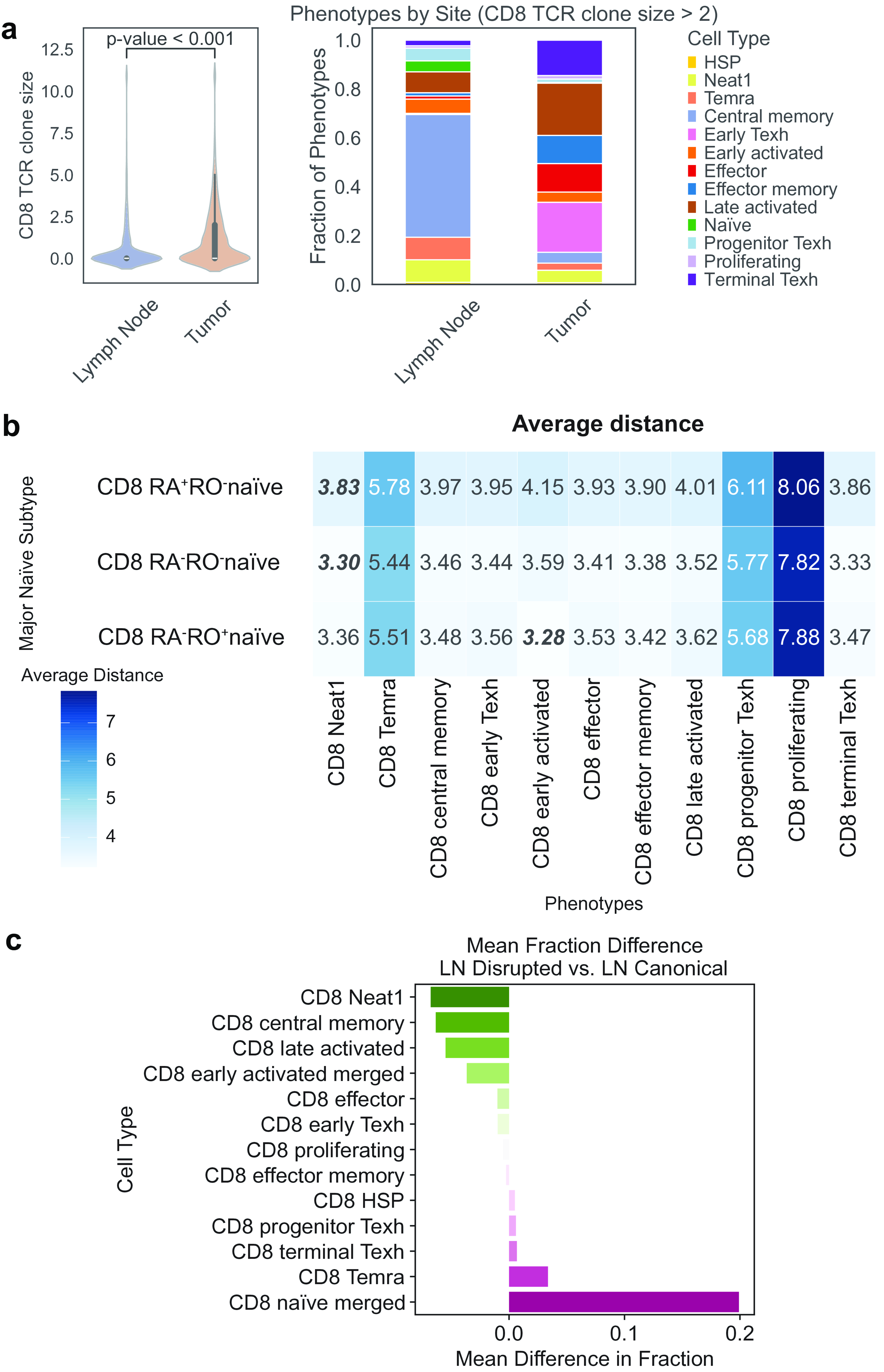

### Extended Figure 5

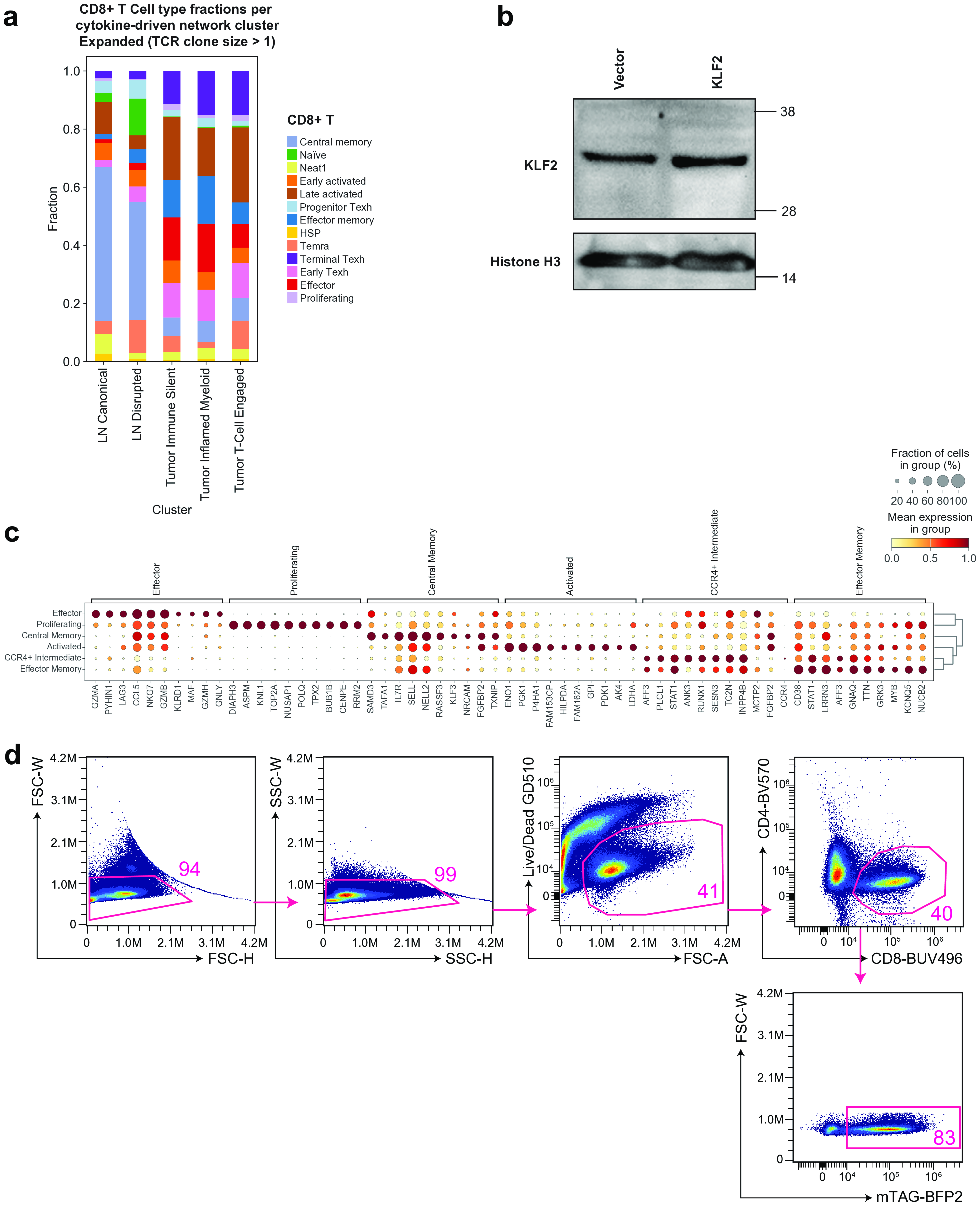

Extended Figure 6

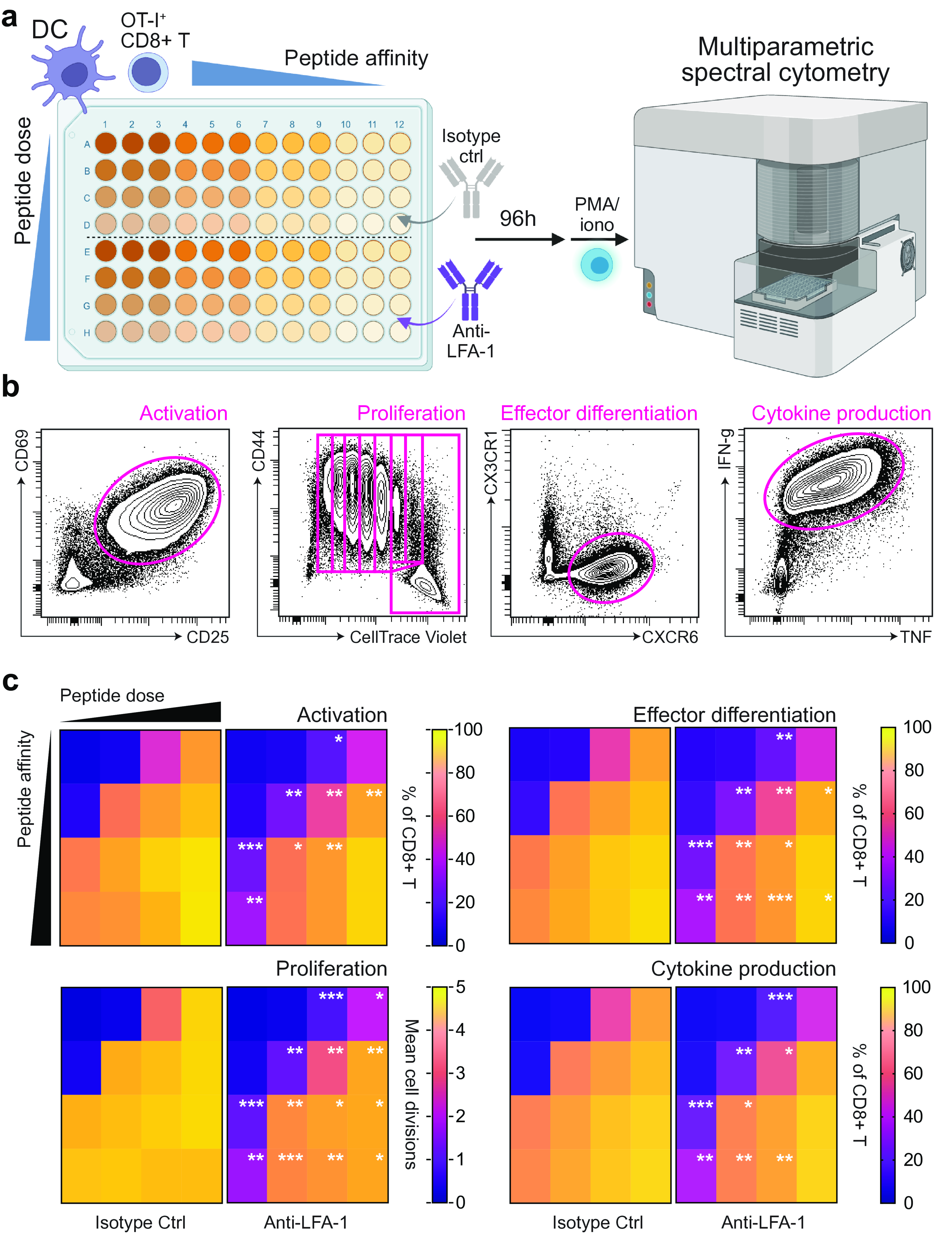

### Extended Figure 7

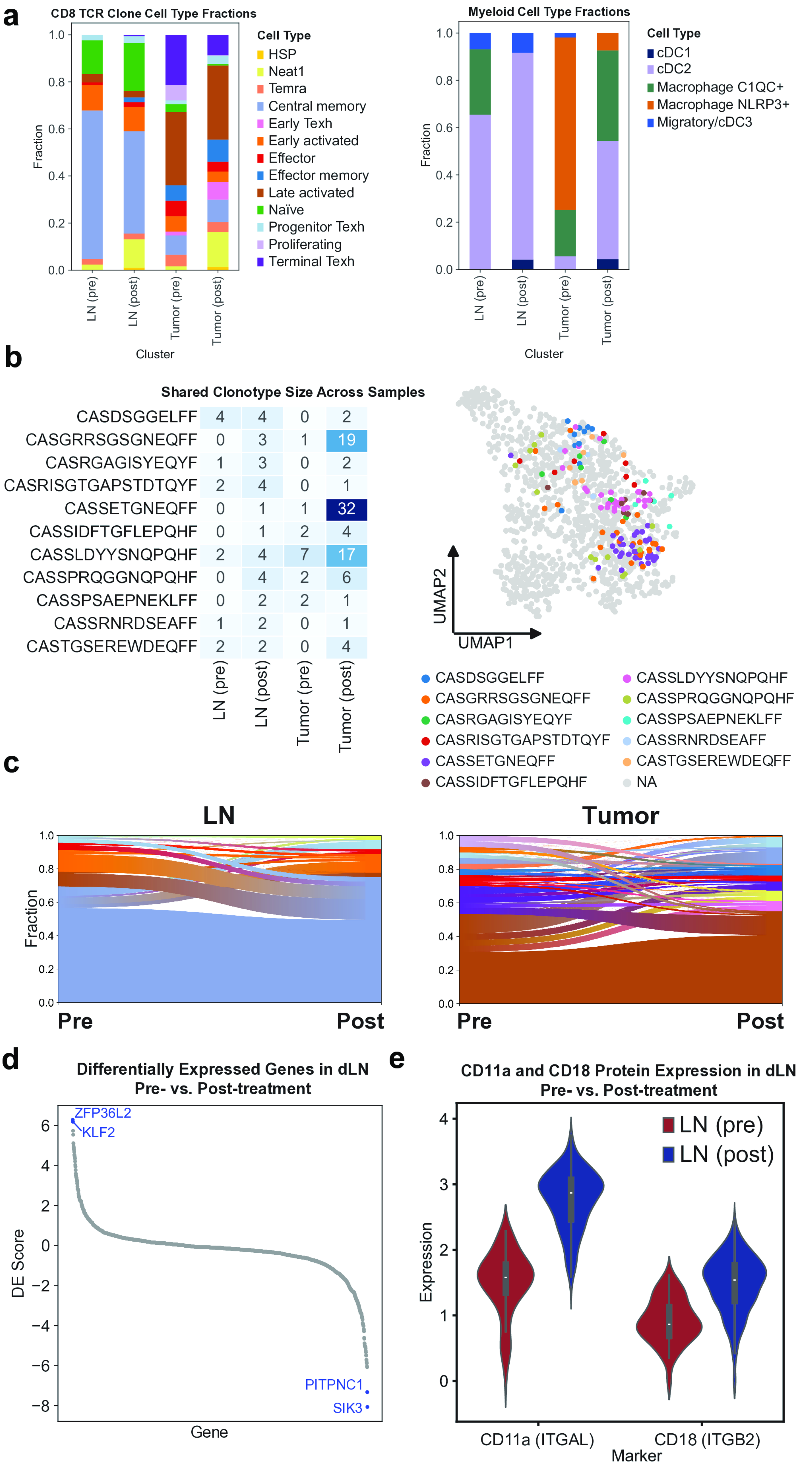
